# Ancestral genomic stasis and coordinated genome erosion underlies asymmetric acarine diversification

**DOI:** 10.64898/2026.09.16.752239

**Authors:** Arjun Cherukutty, Pratik Khopkar, Shreya Salunkhe, Dibyo Mazumder, Siddharth Kulkarni

## Abstract

Mites and ticks (Acari) constitute one of the most species-rich and ecologically diverse arachnid radiations, yet their interrelationships and genomic mechanisms underlying their diversification remain poorly resolved. Here, we reconstructed the evolutionary relationships and trajectory of Parasitiformes using phylogenetics and comparative genomics to test the phylogenetic signal in rare genomic changes. Maximum likelihood and macrosynteny analysis revealed multiple shared chromosomal fusion-with-mixing events that unite Opilioacaridae and Ixodida as sister taxa, with Mesostigmata as a derived sister clade—overturning the prior basal placement of Opilioacaridae as a slowly evolving “living fossil” lineage. We show that major parasitiform lineages diversified through asymmetric, large-scale genome and regulatory rewiring. Opilioacaridae genome retains ancestral chelicerate genomic features, including an intact, contiguous Hox gene cluster, high repetitive-element content, and a conserved microRNA repertoire, all shared with Ixodida. By contrast, the hyperdiverse Mesostigmata shows marked genome compaction, with shortened introns, disrupted and fragmented Hox clusters, and widespread loss of microRNA families, despite retention of developmentally essential miRNAs. Comparative analysis of gene-family evolution further identifies lineage-specific expansions in Ixodida associated with blood feeding and immune evasion, alongside independent, convergent expansion of cuticle-protein gene families in ectoparasitic Mesostigmata. These findings demonstrate that contrasting modes of genome evolution—long-term structural stasis versus rapid compaction—together with convergent gene-family expansion linked to independent origins of parasitism, can coexist within a single arachnid radiation, offering new insight into the molecular basis of developmental regulatory evolution and adaptive diversification in Acari.

## INTRODUCTION

Acari comprises one of the most diverse radiations of Chelicerata, occupying nearly every terrestrial and many aquatic ecosystems and exhibiting extraordinary variation in morphology, ecology, and life-history strategy^1^ spanning free-living predators, scavengers, fungivores, herbivores, and numerous independent origins of parasitism^1,2^. The evolutionary history of Acari remains difficult to reconstruct because major lineages diverged deep in geological time and exhibit elevated rates of molecular evolution, extensive genome rearrangement, and extreme differences in body-plan organization^2–7^. Consequently, relationships among major acarine lineages have remained contentious despite decades of morphological and phylogenomic investigations (Fig. 1g).

**Figure 1.**
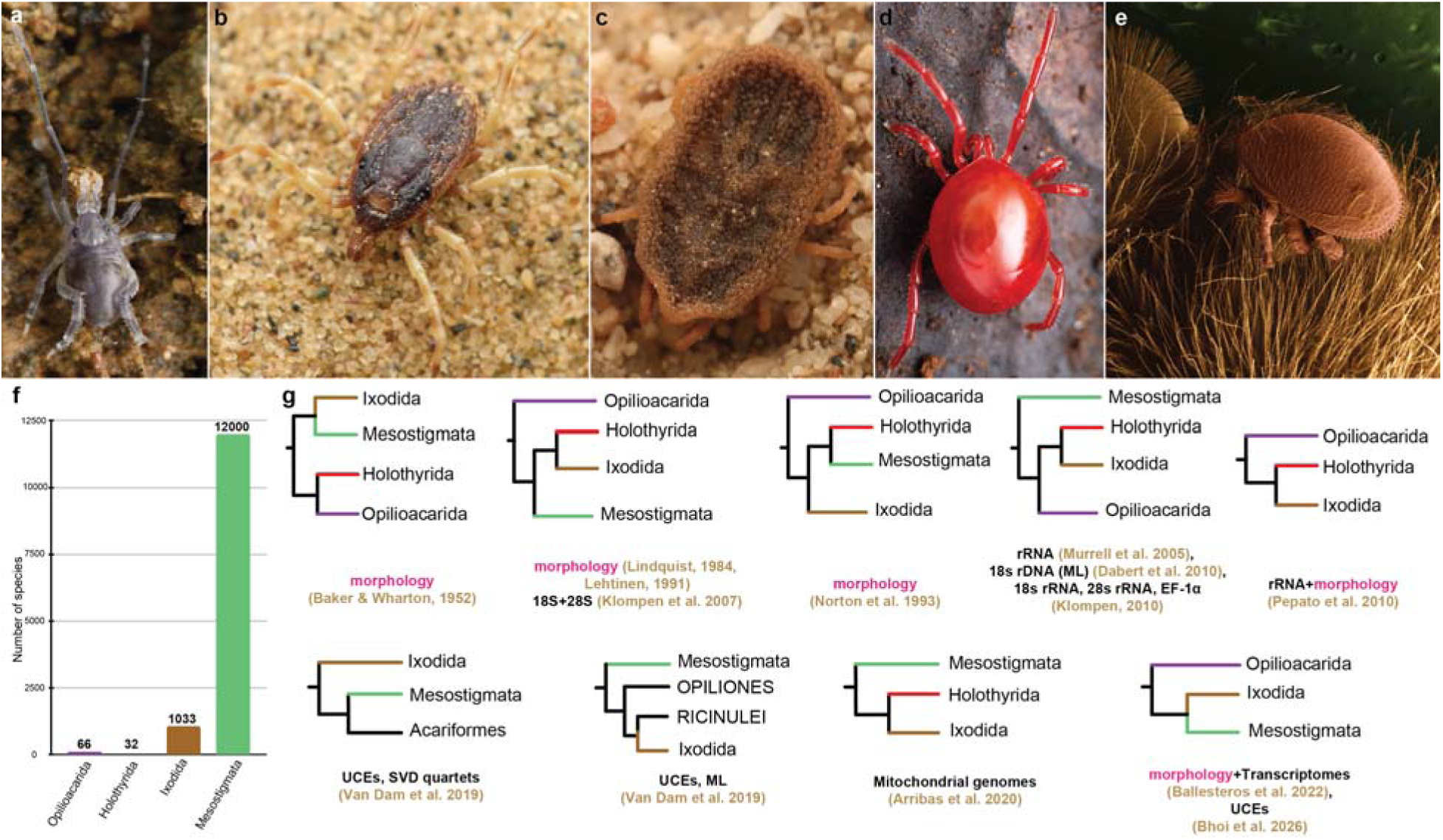
Morphological diversity and competing phylogenetic hypotheses for interrelationships of Parasitiformes. **a—e** Representative members of the four extant parasitiform orders, illustrating the striking disparity in body form, ecology, and degree of morphological specialization within the group. **a** Opilioacarida; **b** Ixodida (Ixodidae, hard ticks); **c** Ixodida (Argasidae, soft ticks); **d** Holothyrida; **e** Mesostigmata (bee mite). Opilioacarids retain several plesiomorphic arachnid features, including visible segmentation and elaborate sensory appendages, whereas ixodid ticks, holothyrids, and mesostigmatans exhibit varying degrees of morphological specialization associated with parasitic, predatory, or detritivorous lifestyles. **f** Asymmetry in described count of species estimated from^8–11^. **g** Summarized alternative phylogenetic hypotheses proposed for relationships among the major parasitiform lineages inferred from different data classes of evidence from early morphology-based studies to genome scale data. Together, these alternative hypotheses highlight the persistent uncertainty surrounding deep parasitiform relationships and require the exploration of characters such as rare genomic changes and macrosynteny as complementary sources of phylogenetic evidence. (Figs. 1a, b © Apurv Jadhav; c © Benjamin Burgunder/Wikimedia Commons; d © Damien Brouste/iNaturalist; e © Eric Erbe, Christopher Pooley: USDA, ARS, EMU/Wikimedia Commons).

Within Acari, Parasitiformes comprises four traditionally recognized orders: Opilioacarida, Ixodida, Holothyrida, and Mesostigmata (Fig. 1a-e)^13–15^. Ixodida includes both soft and hard ticks that exhibit obligate hematophagy, whereas Mesostigmata comprises a diverse assemblage of parasitic and free-living mites. In contrast, Opilioacarida and Holothyrida are known to retain scavenging feeding habits, consuming solid and fluid food resources, respectively^16^. Opilioacarida retains several morphological features considered ancestral for mites, including externally visible opisthosomal segmentation, well-developed eyes, elongated appendages, and complex sensory structures^17^. In contrast, most derived acarine lineages exhibit varying degrees of body-plan attrition, segmental reduction, sensory specialization, and ecological adaptation.

Some phylogenies initially recovered the monophyly of Acari^5,18^, but this result is destabilized by the addition of basally branching groups of Parasitiformes (e.g., Opilioacaridae, the putative sister group to the remaining parasitiforms) or is not supported in sensitivity analyses incorporating taxon permutation, suggesting that Acari monophyly is a long branch attraction artifact^19–22^. Consequently, Opilioacarida have long been viewed as a key lineage for understanding the ancestral condition of Parasitiformes and the evolutionary transitions that generated the remarkable diversity of modern mites and ticks^17,21^.

Despite their evolutionary significance, high quality genomic resources for Opilioacarida remain virtually absent, except a recent short-read genome^3^. Existing acarine genomic datasets are heavily prioritized toward medically important ticks and agriculturally significant mites^23–25^. The absence of a high contiguity genome for Opilioacarida, has limited our ability to reconstruct ancestral genomic states and identify the genomic changes associated with major ecological and morphological transitions.

Rcent advances in comparative genomics have facilitated exploration of rare genomic changes, including whole-genome duplications, transposable-element insertions, gene-order rearrangements, and chromosomal fusion events as phylogenetic characters because they are unlikely to arise convergently^26–30^. Conservation of ancestral linkage groups and chromosomal fusion-with-mixing events have emerged as a robust framework for resolving recalcitrant relationships across Metazoa^31,32^. These shared linkages preserve evidence of common ancestry over deep evolutionary timescales that often obscure sequence-based signals in a phylogenetic framework^33^. Whether such genomic architectural characters can clarify relationships within internal groups of Chelicerata, particularly the hyperdiverse Acari remains underexplored^3,34,35^.

Developmental genome architecture provides a complementary perspective on the evolution of acarine body plans. Hox genes are central regulators of anterior–posterior patterning and segment identity across Bilateria, and their genomic organization is frequently associated with developmental complexity and body-plan evolution^36^. Many arthropods retain relatively conserved Hox clusters^37,38^ while some acarine genomes exhibit extensive Hox-cluster fragmentation, gene loss, and rearrangement^24,39^. These changes have been hypothesized to result in the reduction of the arthropod segmentation and the evolution of highly derived body plans^40,41^. However, the ancestral organization of Hox genes remains uncertain owed to the absence of an Opilioacarid genome.

MicroRNAs (miRNAs), the non-coding regulatory elements, are highly conserved post-transcriptional regulators that influence developmental patterning, neural differentiation, sensory organ development, and tissue specification^42^. Considering miRNA families are rarely regained once lost, patterns of miRNA retention and loss can provide valuable insights into both phylogenetic history and the evolution of developmental complexity^43^.

Acari encompasses some of the most extreme examples of genome expansion and reduction among arthropods^44,45^. Tick genomes can exceed several gigabases, characterized by extensive transposable-element accumulation, whereas many derived mite lineages possess compact genomes ^3,46^. Whether these contrasting genomic states reflect independent evolutionary trajectories or derive from a common ancestral toolkit remains unclear.

We provide a genomic reconstruction of the ancestral parasitiform condition and demonstrate how large-scale and coordinated changes in chromosome organization, developmental regulation, repeat content, and gene architecture contributed to the evolution of extant mite and tick diversity. By integrating comparative analyses of chromosome-scale synteny and other data classes representing rare genomic changes across Opilioacarida using a *de novo* genome, Ixodida, and Mesostigmata, we address the evolutionary relationships among Parasitiformes, functional sources of asymmetric genome architectures, gene family turnovers and genetic toolkit for parasitism.

## RESULTS

### Conserved chromosome architecture supports an Opilioacarida–Ixodida sister-group relationship

#### Genome stats and assembly strategy

Oxford Nanopore sequencing using high molecular weight DNA from 10 individuals yielded raw data of 253.8 Gbp covering 50.9 M reads with an N50 of 7,952 bp. Assembly using hifiasm, followed by purging haplotigs and polishing using purge dups and Racon respectively, recovered a final genome size of 2.93 Gb (N50=2.08 Mb) and a completeness of 94.5% with 86.9% single copy orthologs and 6.24% duplications using Arthropoda ODB version 12 (Supplementary Fig. 2.1). Phylogenomic analyses using Posterior Mean Site Frequency (PMSF) models (C20, C60) of 155 single copy orthologs across Parasitiformes and two acariform outgroups inferred using OrthoFinder recovered Mesostigmata as a sister group to Opilioacarida and Ixodida clade.

### Shared chromosomal fusion-with-mixing events

To investigate conserved phylogenetic signals in macrosynteny as a rare genomic change, we compared the newly assembled opilioacarid genome with chromosome-level genomes from Ixodida (n = 5) and Mesostigmata (n = 3). A custom Parasitiformes Linkage Group (ParaLG) framework using chromosomal assemblies from *Argas vulgaris* (A), *Ixodes scapularis* (I), *Neoseiulus longispinosus* (N) and *Varroa destructor* (V) (Supplementary File 2.1). Multiple ancestral linkage group (ALG) revealed fusion-with-mixing events shared exclusively between Opilioacarida and Ixodida, including AIN11×AIN14×AIN23, AIN1×AIN13×AIV2, AIN16×AIN18, AIN19×AIN3, AIN21×AIN25×AIN12, AIN20×AIN9 and AIN10×AIN15×AIV25 (Fig. 2a). In contrast, Mesostigmata placed these ancestral linkages on separate chromosomal units mixed with other linkage groups (Fig. 2b, Supplementary Fig. 1.1), except for a single fusion event involving AIN19×AIN3 in *Varroa destructor*. This sharing of scores of exclusive fusion-with-mixing events between Opilioacarida and Ixodida constitutes syntenic synapomorphies supporting their sister-group relationship corroborating our phylogenetic results.

**Figure 2.**
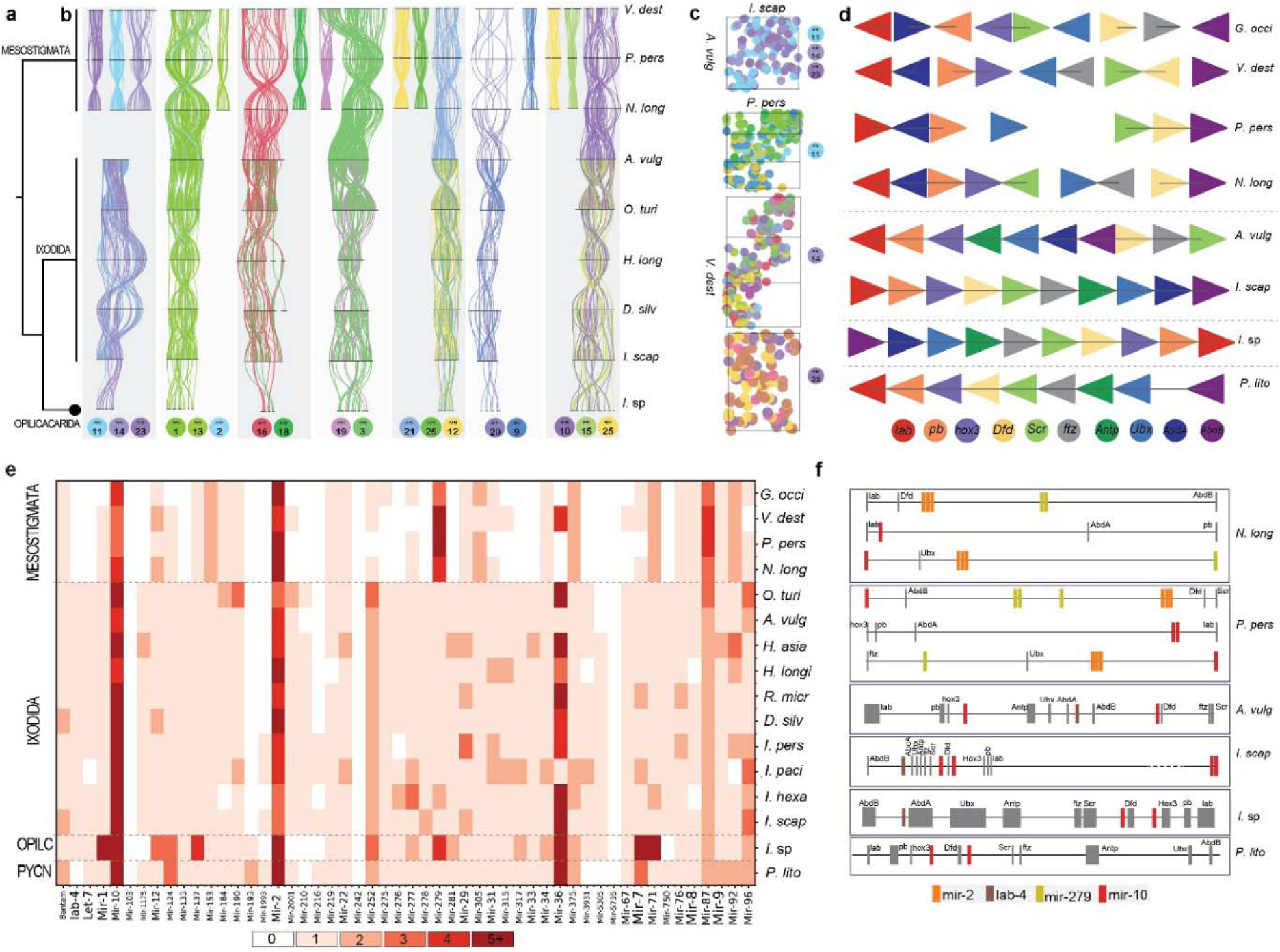
Conservation of ancestral linkage groups, Hox-cluster organization, and microRNA evolution across Parasitiformes. **a** Phylogenetic topology showing the relationship between Opilioacarida, Ixodida and Mesostigmata using macrosynteny reveals Opilioacarida as the sister group of Ixodida. **b**, Macrosynteny analysis comparing highly contiguous assemblies of Parasitiformes using reconstructed Parasitiformes ancestral arachnid linkage groups (ParaLGs). Colored ribbons indicate orthologous genes assigned to each ancestral linkage group, demonstrating large-scale chromosomal conservation and rearrangement. Opilioacaridae and Ixodida retain extensive ancestral linkage-group structure with multiple shared fusion with mixing events, whereas Mesostigmata exhibit substantially greater chromosomal reorganization. Numbers below each linkage group indicate the inferred ancestral ParaLG identity. **c**, Oxford dot plots illustrating chromosomal fusion-with-mixing events as an irreversible rare genomic change among representative Ixodida (top) and Mesostigmata (bottom) genomes. Orthologous genes are plotted according to chromosomal position in pairwise genome comparisons with ParaLG. A shared fusion involving ancestral linkage groups ALG11, ALG14, and ALG23 is evident within Ixodida, whereas these linkage groups remain separated in Mesostigmata. **d**, Microsynteny of Hox genes across Parasitiformes and the outgroup Pycnogonida. Colored triangles denote Hox genes, with orientation indicating transcriptional direction. Opilioacarida and Ixodida retain a largely contiguous and collinear Hox cluster comparable to the inferred ancestral chelicerate condition, whereas Mesostigmata shows increased fragmentation, rearrangement, and dispersion of Hox genes across genomic regions. These patterns indicate disruption of ancestral developmental gene organization during parasitiform evolution. **e**, Heatmap of conserved microRNA family copy numbers across Parasitiformes and the outgroup pycnogonid. Color intensity corresponds to microRNA copy number. Opilioacarida and Ixodida retain a comparatively rich ancestral microRNA repertoire, whereas Mesostigmata show extensive losses and copy-number reductions, consistent with broader patterns of genome streamlining and regulatory compaction. Conserved retention of core families, including mir-2 and mir-10, is evident across all sampled taxa. **f**, Genomic organization of Hox genes and associated microRNAs. Hox genes are shown as gray boxes and conserved microRNAs as colored boxes. Opilioacarida and Ixodida preserve ancestral associations between developmental regulators and non-coding regulatory elements similar to the pycnogonids, whereas derived lineages exhibit extensive genomic rewiring, including altered spacing, rearrangements, and loss of ancestral syntenic relationships. The parallel disruption of Hox organization and microRNA architecture supports a model in which chromosomal reorganization was accompanied by substantial rewiring of developmental regulatory networks during the diversification of Parasitiformes.

### Conservation of developmental genomic architecture and no evidence for ancient whole-genome duplication

Ancient whole-genome duplication (WGD) events, one round in Arachnopulmonata, three rounds in Xiphosura have been proposed using copy number of hox genes, along with a conjecture on a potential round in Acari. However, neither duplicate Hox clusters nor large-scale duplicated chromosomal blocks were detected in Opilioacarida, Ixodida or Mesostigmata (Fig. 2d Supplementary Fig. 1.3a-j), corroborating previous findings about lack of WGD in Acari^34,35^.

#### Retention of a contiguous Hox cluster

Hox gene order is known to be highly conserved across Arthropoda. We screened representative chelicerate genomes for Hox cluster organization across Parasitiformes and the pycnogonid *P.litorale* as an outgroup. In the opilioacarid genome, all ten canonical Hox genes were recovered within a single cluster spanning 11.16 Mbp (Fig. 2c). This conserved order is similar to the organization in Ixodida and Pycnogonida.

The retention of a contiguous Hox cluster in Opilioacarida mirrors the preservation of externally visible opisthosomal segmentation in living opilioacarids. In contrast, derived parasitiform mites exhibit extensive fragmentation of the Hox cluster, with Hox genes distributed across multiple chromosomes. Similarly, Mesostigmata shows substantial disruption of canonical Hox gene order, accompanied by lineage-specific gene losses. Notably, all sampled mesostigmatans lack *Antennapedia*, whereas *Phytoseiulus persimilis* additionally lacks *hox3* and *fushi tarazu* (Fig. 2c).

Differences in gene orientation among taxa likely reflect inversion and translocation events following chromosomal rearrangement. Together, these patterns indicate long-term retention of ancestral Hox architecture in Opilioacarida and Ixodida, whereas Mesostigmata has experienced extensive lineage specific developmental gene reorganization.

#### Conserved microRNA repertoires reveal retention of ancestral regulatory complexity

MicroRNAs (miRNAs) are highly conserved regulatory elements that play essential roles in developmental and physiological processes, including regulation of Hox genes. We identified a broadly conserved core of ancient bilaterian and arthropod miRNA families, including Mir-1, Let-7, Mir-10, Mir-9, Mir-279, Mir-8 and Bantam, across Acari (Fig. 2d; Supplementary File 2.2). Opilioacarida and Ixodida possess comparatively large and intact miRNA repertoires, retaining 336 and 94 miRNA copies, respectively, compared with only 46 copies in Mesostigmata (Supplementary File 2.2). Of the 52 miRNA families screened using MirMachine, 15 families (28%) were completely absent from all sampled mesostigmatan genomes.

Several miRNA families exhibited lineage-specific patterns of retention and loss. Mir-2 and Mir-10 showed high copy numbers across all sampled Parasitiformes, including the pycnogonid outgroup, indicating deep evolutionary conservation. In contrast, Mir-36 displayed extensive copy-number reduction in Mesostigmata, with four copies retained in *Varroa destructor* and absent in all remaining sampled mesostigmatan taxa.

Opilioacarida retained elevated copy numbers of several ancestral miRNA families, including Mir-12, Mir-1, Mir-2, Mir-279, Mir-71, Mir-7 and Mir-36. Multiple copies of Mir-279 and Mir-71 are retained in Opilioacarida and Mesostigmata, whereas Ixodida generally retained only single copies. Conversely, Mir-153, Mir-87, Mir-305 and Mir-92 exhibited higher copy numbers in Mesostigmata than in Opilioacarida and Ixodida.

Together, these results suggest that the acarine ancestor possessed a relatively complex regulatory toolkit that has been differentially retained across descendant lineages. The extensive losses observed in Mesostigmata are consistent with secondary attrition accompanying genome restructuring.

#### Conservation and fragmentation of Hox-associated microRNAs

To investigate the evolution of developmental regulatory architecture, we mapped miRNA locations relative to Hox genes. Of 52 miRNA families examined, eleven were physically associated with Hox clusters (Fig. 2e; Supplementary File 2.8.

Canonical_HOX_miRNAs.tsv). Canonical Hox-linked miRNAs, including Mir-10, Mir-2, Mir-279 and Iab-4, were identified in several taxa. Mir-10 exhibited particularly strong positional conservation, occurring between *hox3*, *Deformed* and *Sex combs reduced* in Opilioacarida and Ixodida. This arrangement likely represents the ancestral state for Chelicerata.

Iab-4 was absent from both Mesostigmata and Pycnogonida, indicating either repeated losses or convergent evolutionary reduction. Interestingly, in Ixodida and Opilioacarida where Iab-4 is present, they are associated with Abd-A which bear the abdominal segments. However, Abd-A too is absent in many Mesostigmata and Pycnogonida that have reduced posterior segments. Multiple copies of Mir-2 and Mir-279 showed Hox-cluster association in Mesostigmata but not in Opilioacarida, Ixodida or Pycnogonida (Fig. 2e).

Overall, Opilioacarida and Ixodida retain canonical Hox-linked miRNAs within conserved genomic neighbourhoods, whereas mesostigmatid mites exhibit extensive fragmentation of both Hox architecture and associated regulatory elements. These observations are consistent with previous suggestions that disruption of Hox-cluster organization may influence segmentation dynamics through altered chromatin architecture and Hox-gene regulation.

### Genome miniaturization and restructuring in Mesostigmata

Genome sizes varied substantially across Parasitiformes (Fig. 3a) (Supplementary Table 2.1). Opilioacarida and Ixodida possessed comparatively large genomes exceeding 1 Gb, whereas mesostigmatan genomes were generally less than 500 Mb. Among ixodids, *Rhipicephalus microplus* possessed the largest genome, exceeding 3 Gb. These results indicate that extensive genome reduction has occurred independently within mesostigmatan lineages. GC content ranged between 45-50% within Ixodida, however, it was above 50% within Mesostigmata (except *Varroa destructor* which was ca. 40%) and ca. 37% in Opilioacarida (Fig. 3a).

**Figure 3.**
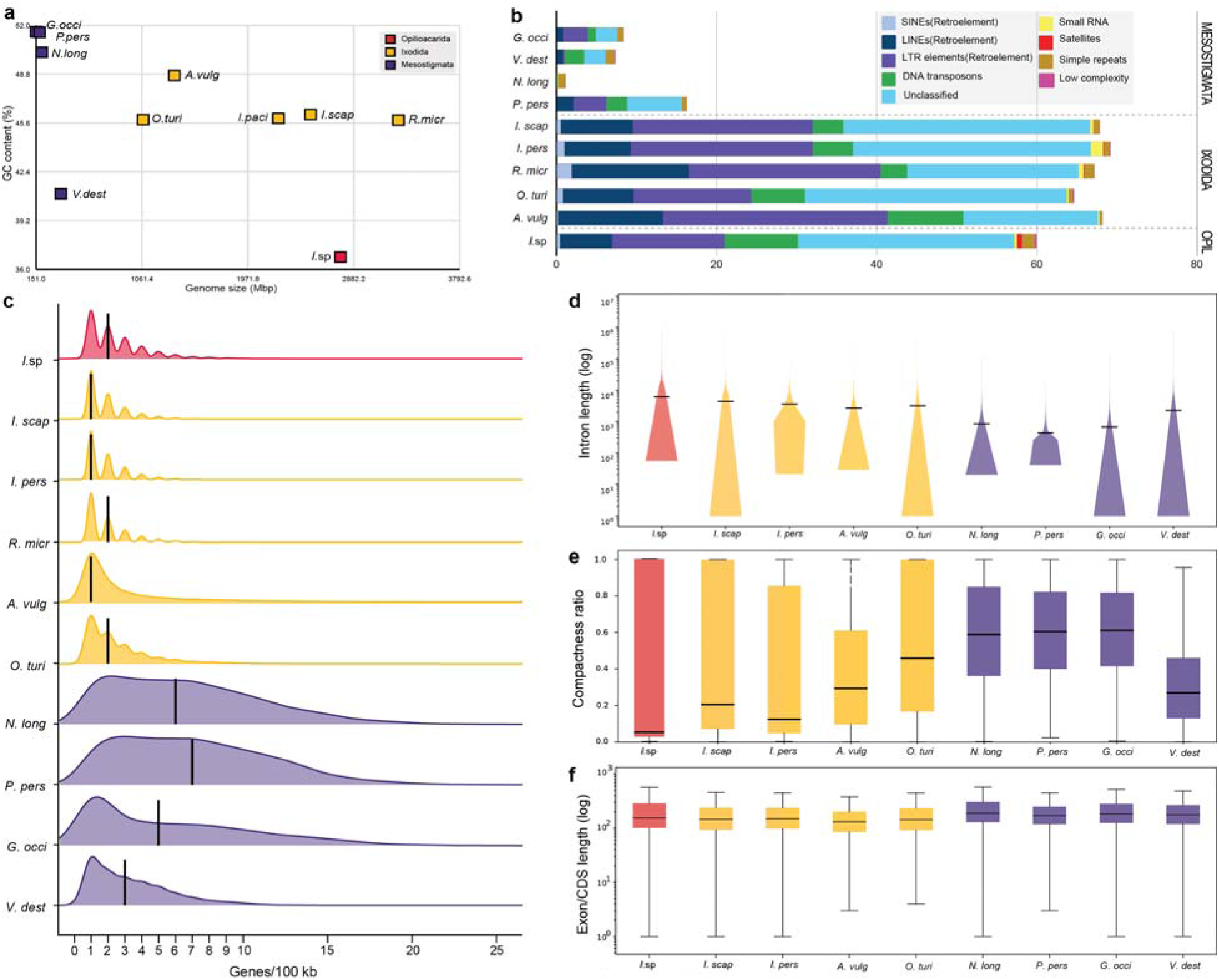
Contrasting trajectories of genome expansion and compaction across Parasitiformes. **a**, Relationship between genome size and GC content across representative Parasitiformes. Opilioacarida occupy an intermediate position characterized by a large genome and relatively low GC content, whereas ixodid ticks exhibit some of the largest genomes sampled. Mesostigmatan genomes have reduced genome sizes and variable GC content, consistent with lineage-specific genome streamlining. **b**, Composition of repetitive elements across parasitiform genomes. Stacked bars show the proportional contribution of major repeat classes relative to parasitiform genome sizes. Ixodid genomes are characterized by extensive repeat accumulation, particularly retrotransposons and unclassified repetitive DNA, whereas Mesostigmata generally exhibit reduced repeat content. The Opilioacaridae genome retains a repeat-rich architecture more similar to ixodids than to miniaturized mesostigmatan genomes, supporting the possibility that repeat-rich genomes represent the ancestral parasitiform condition. **c**, Ridgeline density plots showing the distribution of local gene density per non-overlapping 100 kb genomic window across representative Parasitiform genomes. Vertical black bars indicate the median gene density for each species. Opilioacarida exhibits a broad distribution with low median gene density. Ixodida displays intermediate distributions dominated by low-to-moderate gene densities, reflecting repeat-rich genomes with relatively conserved gene spacing. In contrast, Mesostigmata shows markedly right-shifted distributions and substantially higher median gene densities, indicative of extensive genome compaction. **d**, Distribution of intron lengths across Parasitiformes. Violin plots summarize the variation in intron size for each species. Ixodid and opilioacarid genomes retain broad intron-length distributions with numerous long introns, whereas mesostigmatan genomes are enriched for shorter introns, reflecting extensive genome compaction. **e**, Gene compactness across Parasitiformes, measured as the proportion of a transcript’s genomic span occupied by coding or exonic sequence. Values approaching 1 indicate highly compact genes with limited intronic sequence, whereas lower values reflect intron-rich gene structures. Mesostigmata exhibit significantly higher gene compactness than Opilioacaridae and Ixodida, consistent with pervasive genome streamlining through intron reduction and contraction of non-coding regions. **f**, Exon and coding-sequence (CDS) length distributions across parasitiform genomes. Boxplots summarize variation in exon/CDS length among species. In contrast to the pronounced reduction in intron size and gene span observed in compact genomes, exon and CDS lengths remain comparatively conserved across lineages.

#### Repeat composition differs markedly among major lineages

Repeat elements constituted a substantial fraction of all examined genomes but varied considerably among lineages (Fig. 3b). Mesostigmata contained markedly lower repeat content than either Opilioacarida or Ixodida. Retroelements, including LINEs, SINEs and LTR elements, together with unclassified repeats, accounted for the largest proportion of repetitive DNA (Supplementary Table 2.2). Simple repeats were present across all taxa.

#### Gene density increases with genome compaction in derived Parasitiformes

Genome-wide gene density distributions differed markedly among parasitiform lineages (Fig. 3c). Opilioacaridae and ixodids exhibited low median gene densities with broad distributions, consistent with large genomes containing extensive intronic and intergenic regions. In contrast, mesostigmatid genomes displayed substantially higher gene densities and narrower distributions, reflecting extensive genome compaction (Supplementary File 2.3).

### Intron size distributions reveal conserved gene architecture in Opilioacarida and Ixodida

Comparative analysis of intron size distributions revealed marked differences among major parasitiform lineages (Fig. 3d). The opilioacarid genome exhibited a broad distribution of intron lengths, with median intron sizes comparable to those observed in ixodid genomes. Likewise, ixodids displayed consistently large introns and extended distribution tails, indicating intron-rich gene architectures. In contrast, all sampled mesostigmatan genomes exhibited substantially shorter introns and reduced frequencies of extremely large introns (Supplementary File 2.4). An exception was *Varroa destructor*, which possessed intron lengths approaching those observed in Ixodida. The similarity of intron size distributions between Opilioacarida and Ixodida is consistent with the sister-group relationship recovered in phylogenomic and synteny analyses. These results suggest that intron-rich gene architectures represent the ancestral condition for Parasitiformes and that Mesostigmata underwent secondary intron reduction.

### Gene compactness reveals extensive genomic erosion in Mesostigmata

Gene compactness, defined as the ratio of total exon length to total gene span, revealed substantial differences among lineages (Fig. 3e). Ixodida displayed considerable variation in compactness values, reflecting heterogeneity in intron content and genome size (Supplementary File 2.5). Mesostigmata exhibited more uniform compactness ratios indicative of streamlined gene structures with *Varroa destructor* as an outlier having relatively lower compactness ratio comparable to Ixodida.

Opilioacarida possessed low compactness values consistent with intron-rich genes and expansive non-coding regions. These findings further support the hypothesis that mesostigmatan genomes have undergone extensive secondary compaction.

### Coding sequence lengths remain comparatively conserved

Despite substantial differences in genome size, coding sequence (CDS) lengths were comparatively conserved across Parasitiformes (Fig. 3f). Ixodida exhibited greater variation in CDS lengths than Mesostigmata, whereas Opilioacarida showed CDS lengths comparable to those observed in ticks. The relative stability of CDS lengths suggests that genome size differences primarily reflect variation in non-coding regions rather than extensive changes in protein-coding content (Supplementary File 2.6).

### Repeat landscape evolution across Parasitiformes

#### Transposable-element age distributions reveal contrasting evolutionary histories

To investigate the timing of transposable-element activity, we analysed sequence divergence between TE copies and their reconstructed consensus sequences. Low divergence values indicate recent insertions and high divergence values indicate ancient insertions. *Ixodida* exhibited pronounced peaks at 3–8% divergence, indicating substantial recent TE activity and showed broader distributions spanning approximately 25–35% divergence, consistent with multiple episodes of TE expansion over extended evolutionary periods. In contrast, the opilioacarid genome exhibited a broader divergence peak spanning 5–30%, suggesting that their major TE expansions occurred earlier and that recent activity has been relatively limited. The mesostigmate *Phytoseiulus persimilis* displayed a distinct peak near 25% divergence, indicating an ancient burst of transposable-element activity followed by prolonged quiescence.

*Neoseiulus* exhibited extremely low repeat abundance and lacked prominent divergence peaks, consistent with efficient long-term TE suppression or removal (Figs. 4a-c; Supplementary File 2.7). Collectively, these findings identify Opilioacarida as retaining an intermediate and relatively stable transposable-element landscape. In contrast, tick genomes have undergone repeat-driven expansion, whereas mesostigmatan lineages exhibit signatures of repeat loss and genome miniaturization.

**Figure 4.**
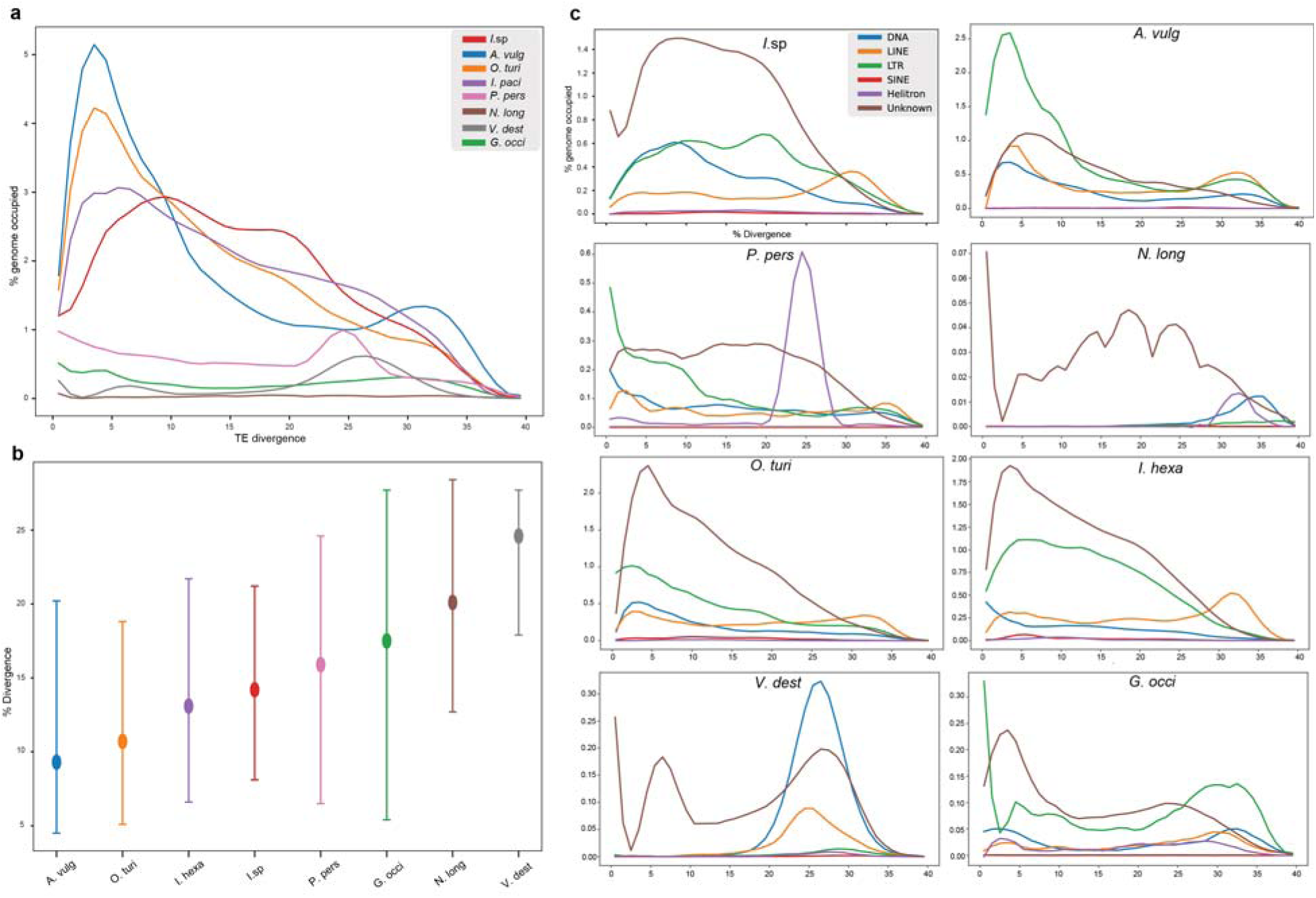
Transposable element (TE) landscapes reveal contrasting histories of genome expansion and streamlining across Parasitiformes. **a**, Repeat landscape profiles of all taxa showing the genomic proportion occupied by transposable elements as a function of sequence divergence from their inferred consensus sequences. Peaks at low divergence indicate recent TE activity, whereas peaks at higher divergence represent older TE expansions. Ixodid genomes exhibit broad TE age distributions consistent with prolonged accumulation and retention of repetitive DNA. In contrast, Mesostigmata display substantially reduced repeat landscapes, indicative of repeat-element depletion and genome streamlining. The opilioacarid genome retains an intermediate but repeat-rich profile, supporting the inference that extensive TE content represents an ancestral parasitiform condition. **b**, Mean sequence divergence of transposable elements across sampled genomes. Points represent average divergence from repeat consensus sequences and error bars denote standard deviation. Lower divergence values indicate relatively recent TE activity, whereas higher values reflect older, more degraded repeat landscapes. Mesostigmatan genomes generally exhibit older and more eroded repeat complements, consistent with long-term repeat loss, while ixodid and opilioacarid genomes retain evidence of more persistent TE accumulation. **c**, Class-specific TE landscapes partitioned by major repeat categories. Ixodid genomes are dominated by multiple TE classes with evidence of repeated expansion events, whereas Mesostigmata show markedly reduced abundance across most categories. The opilioacarid genome retains diverse TE families spanning a broad divergence spectrum, resembling ixodid genomes more closely than compact mesostigmatan genomes. These patterns indicate that genome-size evolution within Parasitiformes has been strongly influenced by lineage-specific differences in TE retention, turnover, and purging.

#### Gene-family turnover accompanies independent transitions to ectoparasitism

Extensive gene-family turnover, with particularly pronounced expansion and contraction across Ixodida was detected (Figs.5 a,b). Among the annotated families using eggnog-mapper, the Opilioacarida+Ixodida ancestral branch showed expansions in acyl-transferase, serpin, sulfatase-associated, carboxypeptidase, chloride-channel and lectin-associated families, whereas Ixodida subsequently expanded sensory, lipid-metabolism and extracellular protease-inhibitor families, including a Kunitz/WAP/Kazal-rich orthogroup (Fig. 5b). Mesostigmata displayed a distinct pattern, with fewer annotated expansions and contractions affecting sensory, detoxification, lipid-metabolic, oxidative-stress, transporter and extracellular-protein families (Fig. 5c, Supplementary Table. 2.4).

**Figure 5.**
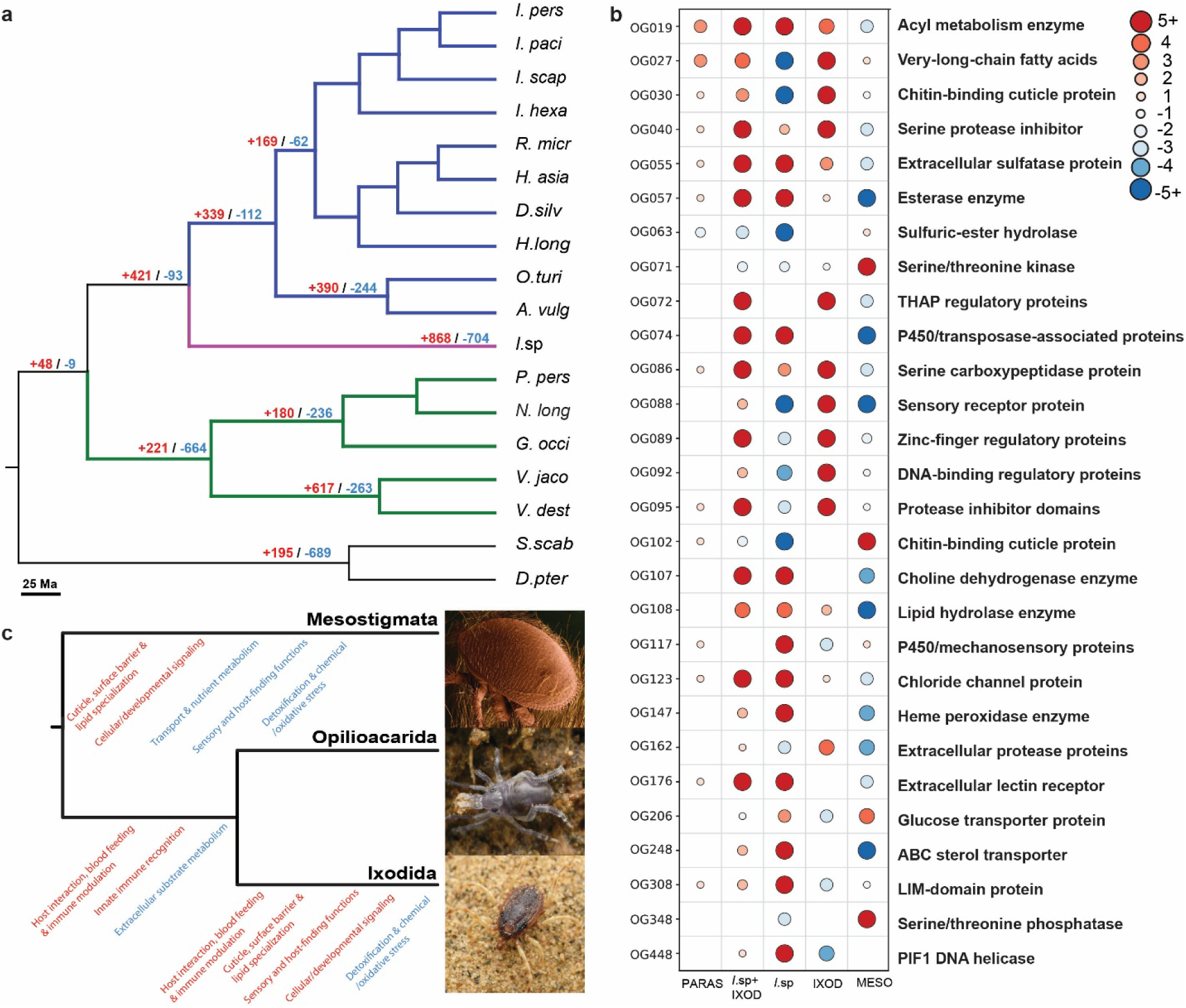
Lineage specific gene family turnover in Parasitiformes. **a.** Gene family expansions (+) and contractions (-) of key nodes in Acari. Note the extensive gains in Opilioacarida (pink)+Ixodida (blue) clade and losses in Mesostigmata crown node (green). Ectoparasite mesostigmatid *Varroa* shows severe expansion, compared to their free-living relatives. **b.** Comparative map of select skewed gene family turnover functions in parasitiforms. Warmer and cooler colors indicate expansions and contractions respectively. Significant patterns include expanded serpins/protease inhibitors and digestive/extracellular proteins supports host manipulation, blood feeding, immune evasion and nutrient acquisition demonstrating key signatures of blood feeding parasitic strategy in Ixodida (OG040, OG055, OG086, OG095, OG162). Cuticle associated gene families expanded in Ixodida and convergently in the mesostigmatid ectoparasite *Varroa* (OG019, OG027, OG030, OG102, OG108). **c.** Select functional groups of gene families expanded (red) and contracted (blue) in major parasitiform lineages. (© Apurv Jadhav; Eric Erbe, Christopher Pooley: USDA, ARS, EMU/Wikimedia Commons).

At the genome-wide, all-family scale, the branches marking the two independent origins of ectoparasitism in this tree. The Ixodida crown stem (339 families expanded:112 contracted) and the *Varroa* stem (617:263) showed a markedly higher expansion-to-contraction ratio than their immediate free-living relatives, Opilioacarida (868:704) and Phytoseiidae (180:236, net contraction) (Fig. 5a). Within a smaller, PFAM/EggNOG-annotated subset of large-effect orthogroups, cuticle-protein genes (Chitin_bind_4) independently expanded on both the Ixodida and Mesostigmata-stem branches while contracting in Opilioacarida, whereas anti-hemostatic (Kunitz/antistasin), immune-lectin (ficolin), and chemosensory (G protein-coupled receptor-GPCR) expansions were tick-specific (Fig. 5c). A large share of the single-orthogroup copy-number swings, especially in Opilioacarida, was attributable to transposon-associated domains rather than conventional gene-family evolution.

## DISCUSSION

The highly contiguous genome assembly of opilioacarid mite provides a long-sought linchpin resource to discern ancestral genomic architecture of Parasitiformes and, more broadly, for understanding the evolutionary processes that generated the remarkable diversity of extant Acari. Previous studies demonstrated the value of Opilioacarida for resolving deep acarine relationships using chromosome-scale genome architecture^3,21^. Here, we show that Opilioacarida also preserves multiple layers of ancestral genomic organization—including macrosynteny, developmental gene architecture, non-coding regulatory repertoires, sensory gene diversity, and transposable element landscapes— that collectively define a genomic baseline from which the derived architectures of modern mites evolved.

Our results reveal a striking evolutionary dichotomy within Parasitiformes. Opilioacarida and Ixodida retain conserved chromosomal organization, large intron-rich genomes, extensive developmental regulatory repertoires, and comparatively stable Hox architecture. In contrast, Mesostigmata exhibit extensive chromosomal reorganization, genome compaction, regulatory attrition, and developmental gene remodeling. These findings suggest that the evolutionary history of Parasitiformes was shaped not by gradual modification of a uniform genomic template, but by fundamentally different trajectories of genome evolution, ranging from long-term architectural conservation to extensive genomic restructuring.

### Conserved genome architecture supports an ancestral Opilioacarida–Ixodida genomic stasis

Deep phylogenetic relationships within Acari have long been difficult to resolve because rapid sequence evolution, lineage-specific rate heterogeneity, and extensive morphological convergence can obscure historical signal in conventional phylogenomic datasets^5,7,8,19^. Increasingly, rare genomic changes have emerged as powerful complementary markers because their probability of independent origin is substantially lower than that of sequence substitutions^26,31^.

Our macrosynteny analyses demonstrate that Opilioacarida and Ixodida share multiple ancestral linkage-group fusion-with-mixing events that are absent from Mesostigmata. Such chromosomal signatures represent complex genomic characters that are unlikely to revert once extensive mixing has occurred^31^. The shared retention of these fusion-derived linkage associations provides independent support for an Opilioacarida–Ixodida relationship previously recovered using 18S rRNA, 28S rRNA and EF-1α^47^. More broadly, the concordance among macrosynteny, Hox organization, intron architecture, and microRNA retention suggests that these lineages have maintained substantial components of the ancestral chelicerate genome over hundreds of millions of years.

This stability of genomic architecture recovering multiple fusion with mixing events robustly supporting a particular relationship seems to be a rare case than a pattern^48^ (Kulkarni et al. *preprint*). For example, even within Arachnida, the sister group of Parasitiformes is contested, due to an ancient rapid radiation in the early Paleozoic, resulting in a soft polytomy shared with Acariformes, Opiliones, Palpigradi, Ricinulei, Solifugae and Xiphosura^7,20^. Macrosyntenic screening of available genomes of some of these orders, support conflicting inter-ordinal relationships by multiple mutually exclusive linkages^35^.

This problem is further obfuscated by the three rounds of whole genome duplication in Xiphosura and one in Arachnopulmonata (spiders, scorpions, pseudoscorpions and pedipalpi)^7,29,34,35^. Notably, we find no evidence supporting an ancient whole-genome duplication in Parasitiformes. Although a possible duplication event has previously been suggested for ticks^49^, neither Hox gene organization nor genome-wide patterns of gene retention support a WGD signal in Opilioacarida or Ixodida. Instead, our results are consistent with previous studies indicating that Parasitiformes retain the plesiomorphic unduplicated genomic condition.

### Genome compaction in Mesostigmata reflects coordinated restructuring of coding and non-coding architecture

Genome sizes vary dramatically across Acari, ranging from highly expanded tick and the opilioacarid genomes exceeding 2 Gb, to some of the smallest arthropod genomes known^44^ (Fig. 3a). The mechanisms underlying this variation remain poorly understood because most comparative analyses have focused either on individual species or specific gene families.

Our results indicate that genome miniaturization in Mesostigmata is not driven by a single process but instead reflects coordinated restructuring across multiple genomic compartments. Compared with Opilioacarida and Ixodida, mesostigmatid genomes exhibit reduced intron lengths, increased gene compactness, diminished repeat content, and extensive chromosomal reorganization. The similarity of intron size distributions between Opilioacarida and Ixodida suggests that intron-rich gene structures represent the ancestral condition for Parasitiformes, whereas Mesostigmata underwent secondary intron reduction and compaction following divergence from the common ancestor.

Importantly, these changes extend beyond protein-coding regions. Repeat landscapes reveal substantial depletion of repetitive DNA in Mesostigmata relative to Opilioacarida and Ixodida. In contrast, the opilioacarid genome retains an intermediate transposable element composition dominated by older DNA transposons and lacks the recent retrotransposon expansions characteristic of several tick genomes^46,50^. Together, these observations suggest that both genome expansion and miniaturization represent derived evolutionary states, whereas the opilioacarid genome more closely approximates the ancestral parasitiform condition.

The transposable element divergence profiles further illuminate these contrasting trajectories. Mesostigmatid genomes show reduced repeat loads, evidence of long-term repeat depletion and ancient TE expansion. Tick genomes exhibit signatures of prolonged or recurrent transposable element activity, whereas Opilioacarida displays predominantly older insertions with relatively limited evidence of recent expansion. The inference of a recent surge of insertion of transposable elements in Ixodida, gradual curve in Opilioacaridae, but ancient expansion in Mesostigmata is intriguing as it defies the traditional notion that Mesostigmata as ‘derived’ and, ixodids and opilioacarids to be primitive. Present fossil record limits Ixodida and Opilioacarida to Late Cretaceous Age^51,52^, (99 Ma), whereas mesostigmatid fossils date back to Early Cretaceous^51^, (∼130 Ma). Our dated phylogeny optimized marginally differing median ages for Ixodida+Opilioacarida clade (212.2 Ma) and Mesostigmata (196.9 Ma) (Supplementary Fig. 3).

The traditional view of Opilioacarida as a primitive lineage is challenged by the possibility that several of its distinctive features are likely derived rather than ancestral^47^. In particular, the absence of abdominal sclerotization may represent a secondary loss, given that reduced or absent sclerotization also occurs in several taxa within Holothyrida and Ixodida. The mid-sternal position of the male genital orifice is shared with Holothyrida, Ixodida, and basal Mesostigmata and therefore does not support a uniquely primitive status for Opilioacarida. Furthermore, features such as multiple stigmata and With’s organ have been proposed as derived specializations rather than retained ancestral conditions^53^. Collectively, these observations along with multiple lines of evidence from genome architecture support the Opilioacarida as a derived lineage within Parasitiformes rather than a basal transitional group. Finally, the observed large-scale gene loss indicates that mesostigmatid lineages have shed significant portions of the ancestral acarine toolkit, likely as an adaptation to specialized predatory or parasitic niches^54,55^.

### Hox cluster integrity links developmental genome architecture to body-plan evolution

One of the most striking findings of this study is the retention of a largely contiguous Hox cluster in Opilioacarida. We recovered all the ten canonical arthropod Hox genes within a single genomic interval spanning 11.16 Mb, preserving a degree of collinearity absent from most derived mite genomes.

Hox genes are central regulators of anterior–posterior patterning and segment identity throughout Bilateria^56,57^. Their physical clustering is often associated with coordinated regulation and conserved developmental patterning^58^. The retention of a contiguous Hox cluster in Opilioacarida mirrors the persistence of externally visible opisthosomal segmentation in the adult body plan, suggesting a correspondence between developmental gene architecture and morphological organization.

In contrast, derived mite lineages exhibit extensive Hox cluster fragmentation. *Metaseiulus occidentalis* represents one of the most extreme examples known among arthropods, with Hox genes dispersed across separate genomic regions rather than maintained in a recognizable cluster^24^. Comparable fragmentation is also evident in other derived acarine genomes. While causality cannot be inferred directly from comparative genomic data, the repeated association between Hox disintegration, reduction of posterior segmentation, and extensive chromosomal reorganization suggests that developmental gene remodeling accompanied major transitions in acarine body-plan evolution.

The evolutionary significance of Hox cluster fragmentation may extend beyond simple gene rearrangement. Pace et al. ^59^ proposed that the loss of abd-A in mites could alter chromatin-level regulation of posterior Hox genes and contribute to developmental truncation of posterior segments. In pycnogonida, the loss of abd-A has been proposed to lead to reduction of posterior tagma^41^. This rewiring of the developmental regulatory networks is also supported by the absence of several PRD, ANTP, TALE, CUT, and zinc-finger homeobox genes in Mesostigmata, contrasted with their retention in Opilioacarida and Ixodida. The preservation of a contiguous Hox cluster in Opilioacarida may represent a retained ancestral developmental state, whereas derived mites evolved alternative regulatory architectures associated with reduced segmentation and body-plan attrition.

### Regulatory evolution mirrors chromosomal restructuring

The microRNA analyses reveal a coordinated pattern of conservation and reduction across Parasitiformes. Opilioacarida and Ixodida retain comparatively large microRNA repertoires and relatively intact genomic neighborhood, whereas Mesostigmata have experienced extensive fragmentation and losses of both microRNA families and copy numbers. Of the 52 microRNA families surveyed, nearly one-third were absent from all sampled mesostigmatid genomes (Fig. 2d).

Several observations suggest that these losses are non-random. Highly conserved developmental families, including mir-2 and mir-10, are retained across Parasitiformes and even in the pycnogonid outgroup. In contrast, more lineage-restricted families exhibit patchy retention and repeated losses. This pattern is consistent with broader observations that deeply conserved microRNAs tend to be maintained because they occupy central positions within developmental regulatory networks^60^. The congruence among chromosomal reshuffling, Hox disruption, and microRNA attrition suggests that genome compaction was accompanied by substantial remodeling of ancestral regulatory networks including reduced posterior segments.

More generally, the data support a model in which the acarine ancestor possessed a relatively complex regulatory toolkit that was differentially retained across descendant lineages. Rather than acquiring novel regulatory complexity, many derived mite groups appear to have evolved through progressive rewiring and loss of ancestral developmental systems. Such convergence suggests that adaptation to specialized ecological niches repeatedly favored the elimination of ancestral genomic complexity rather than the acquisition of novel gene complements.

These findings suggest that the acarine ancestor was not a sensory-reduced organism adapted to cryptic habitats. Instead, it likely possessed a relatively rich visual and chemosensory toolkit that supported active environmental perception. The reduced sensory repertoires observed in many modern mites therefore appear to represent secondary losses associated with ecological specialization, including parasitism, soil dwelling, host dependence, and extreme miniaturization.

The convergent excess of gene-family expansion over contraction at both independent origins of ectoparasitism—ixodids and *Varroa*, relative to their free-living sister lineages, suggests that the shift to an ectoparasitic lifestyle is associated with net gene-family gain. This plausibly reflects shared demands, such as cuticular resilience under host-imposed mechanical and immune stress. It is consistent with the parallel Chitin_bind_4 expansion detected in both lineages. However, this might not a universal signature of parasitism in Acari. The more distantly related parasitic mite *Sarcoptes*, albeit treated as outgroup and therefore poorly sampled, showed the opposite, strongly contraction-dominated pattern. Gene-family dynamics therefore likely track the specific ecological demands of a parasitic strategy, rather than parasitism as a category per se. The tick-specific anti-hemostatic, immune, and chemosensory expansions most plausibly reflect solutions to the distinct challenge of penetrating vertebrate skin and prolonged blood feeding. *Varroa*, the extraoral tissue feeder does not face this problem, which may explain why convergence between the two lineages appears stronger at the level of overall family-turnover balance than at the level of shared homologous gene families. Confirming whether the cuticle-protein signal is specific to *Varroa*, rather than ancestral to all Mesostigmata, remains a necessary next step.

## CONCLUSION

Our findings support a model in which the earliest parasitiform genomes were characterized by chromosomal stability, extensive non-coding architecture, intact developmental gene clusters, rich sensory repertoires, and comparatively complex regulatory systems—features largely retained by Opilioacarida and, to a lesser extent, Ixodida, whereas Mesostigmata experienced extensive chromosomal reorganization, genome compaction, regulatory attrition, and developmental gene remodeling.

Chromosomal architecture and gene-family turnover evolve along partly decoupled trajectories: Ixodida pairs ancestral-like chromosomal stasis with lineage-specific gene-family gain, whereas *Varroa* reverses its clade’s ancestral compaction trend at the branch marking its transition to ectoparasitism. This reframes acarine genome evolution as a balance between long-term architectural conservation and repeated, mechanistically distinct episodes of reductive and expansive change, with Opilioacarida serving as a critical reference for ancestral biology even as a substantial share of its apparent turnover traces to transposable-element activity rather than canonical gene duplication. Future studies integrating chromosome-scale assemblies from the last unsampled parasitiform lineage, Holothyrida, together with finer sampling within Mesostigmata, and spatial transcriptomic and functional genomic approaches, will be essential for testing how these architectural and gene-family changes jointly shaped developmental regulation, sensory function, and host association during the diversification of mites and ticks. By clarifying ancestral genome architecture in a clade whose ancient divergences have been obscured by extensive genomic rewiring, we move closer to resolving the apulmonate soft polytomy within Chelicerata.

The convergent excess of gene-family expansion over contraction at both independent origins of ectoparasitism—ticks and *Varroa*, relative to their free-living sister lineages— suggests that the shift to an ectoparasitic lifestyle is associated with net gene-family gain. This plausibly reflects shared demands, such as cuticular resilience under host-imposed mechanical and immune stress. It is consistent with the parallel *Chitin_bind_4* expansion detected in both lineages. However, this is not a universal signature of parasitism in Acari. The more distantly related parasitic mite *Sarcoptes* showed the opposite, strongly contraction-dominated pattern. Gene-family dynamics therefore likely track the specific ecological demands of a parasitic strategy, rather than parasitism as a category per se.

The tick-specific anti-hemostatic, immune, and chemosensory expansions most plausibly reflect solutions to the distinct challenge of penetrating vertebrate skin and prolonged blood feeding. *Varroa* does not face this problem, which may explain why convergence between the two lineages appears stronger at the level of overall family-turnover balance than at the level of shared homologous gene families. Confirming whether the cuticle-protein signal is specific to *Varroa*, rather than ancestral to all Mesostigmata, remains a necessary next step. Ruling out transposon-driven inflation of the largest orthogroup swings is likewise required. Only then can these patterns be interpreted as confident evidence of convergent molecular adaptation to ectoparasitism.

## Materials and Methods

### Genome sequencing, assembly and annotation

Unfed for a week and flash frozen adult specimens of Opilioacaridae (n=10) were pooled for DNA extraction using the phenol-chloroform method. DNA sample quality was tested with NanoDrop and DNA fragment sizes by agarose gel electrophoresis. Sequencing libraries were prepared using the manufacturer’s protocol and long-read data was generated on the Oxford Nanopore Technology (ONT) Promethion R10.4.1 flow cell. Bases were called from pod5 files using Dorado version 1.1.1+e72f1492 SUP (Super-Accurate)^58^ (available at https://github.com/nanoporetech/dorado) model followed by bam to fastq conversion using bedtools.

Genome was assembled using Hifiasm version 0.25.0-r726c^59^. Assembly quality revealed >10% duplicated BUSCOs (Benchmarking Universal Single-Copy Orthologs) (version 6.0.0 with arthropoda_odb12), significantly more than the other arachnid assemblies. We reason that this could be due to high heterozygosity^60^. To remove false duplications, we employed a second round of haplotig-duplication purging on the opilioacarid genome assembly using purge_dups version 0.0.3 followed by screening to remove potential contamination with Foreign Contamination Screening (FCS-GX)^61^.

Another round of assembly was conducted using FLYe version 2.9.6-b1802^62^ followed by three rounds of polishing using Racon version 1.5.0^63^. Haplotig-duplication were removed using purge_dups for this assembly and further screen for corrections using LongStitch version 1.0.5^64^. The Hifiasm and FLYe assemblies were then merged using Quickmerge version^65^ with its merge wrapper script.

A species-specific repeat library was generated independently for this genome using RepeatModeler2 version 2.0.6^66^. Repetitive elements were subsequently soft-masked using RepeatMasker v.4.1.8^67^. Protein-coding genes were annotated using GALBA version 1.0.11^68–71^ with gene prediction performed using protein homology evidence from UniProt^72^ (https://www.uniprot.org/uniprotkb?query=arachnida; accessed March 2026). This Opilioacaridae genome was used in a comparative framework chromosome-level or highly contiguous assemblies from Ixodida, Mesostigmata, and Pycnogonida as an outgroup for the further analyses.

### Construction of Parasitiformes linkage groups

To identify conserved ancestral linkage associations within Parasitiformes, we developed a custom Parasitiformes Linkage Group (ParaLG) framework using chromosome-scale genomes from *Ixodes scapularis* (Ixodidae), *Argas vulgaris* (Argasidae), *Ornithodoros turicata* (Argasidae), *Varroa destructor* (Varroidae), and *Neoseiulus longispinosus* (Phytoseiidae). Linkage groups were designed and inferred using the Orthology Detection Pipeline (ODP) v.0.3.3^26^. Orthologous genes were identified using a three-way reciprocal-best hits (RBHs) of protein generated with DIAMOND^72^ and OrthoFinder v.2.3.7^73^. The filtered RBHs were grouped using *A.vulgaris* as anchor species to merge and align the three groups.

### Macrosynteny analyses

Oxford dot plots and ribbon plots were generated using ODP to visualize conserved syntenic relationships within Opilioacaridae, Ixodida, Mesostigmata, and outgroup taxa (Supplementary Figure 1.2 a-h). Chromosome annotation files (.chrom) were generated from GFF3 annotations using a custom script (gff_to_chrom.py). Statistical support for linkage-group conservation was evaluated using 1,000,000 randomizations with a false-discovery threshold of α = 0.05.

To identify chromosomal fusion-with-mixing events and other syntenic synapomorphies, linkage-group coordinates were analysed using a custom pipeline 1.better-identify_FwM.py and 2.filter-rbh.py files (https://github.com/kulkarni-lab/GenomeCartographer/tree/main).

### Phylogenomic analyses and orthogroup evolution

Orthologous gene families were identified using OrthoFinder version 3.0.1b1. Protein alignments were generated from single-copy orthologs (SCOs) using MAFFT version 7.525^74^. Maximum-likelihood phylogenies were reconstructed using IQ-TREE version 3.0.1^75^ under the PMSF model with automated model selection and 1,000 ultrafast bootstrap replicates. The resulting topology was used for reconstructing a time calibrated phylogeny using fossil calibrations constrained at nodes (Supplementary file 2.9) using MCMCTree^76,77^. version 4.10.10. Evolutionary dynamics of genome evolution was explored using CAFE5^78^ to map lineage specific expansion and contraction of gene families with a random birth–death process model. Gene family counts were corrected for outliers using base error correction in CAFE5. To account for rate heterogeneity across gene families, we compared CAFE5 Gamma models with increasing numbers of discrete rate categories (k = 1–9) using maximum likelihood. Model fit improved substantially from the single-rate Base model (-lnL = 143,543) to a Gamma model with two categories (-lnL = 135,438), and continued to improve markedly through k = 3 (-lnL = 134,743) and k = 4 (-lnL = 134,613). Beyond k = 4, additional rate categories yielded only marginal gains in likelihood (Δ-lnL < 100 per added category through k = 9), indicating diminishing returns from increased model complexity. We therefore selected the k = 4 Gamma model as it captured the substantial majority of the likelihood improvement over simpler models while avoiding overparameterization associated with higher k values, consistent with recommendations to balance model fit against complexity when the marginal likelihood gain no longer justifies the additional free parameters. Orthogroups with extreme expansions and contractions in key clades (Parasitiformes, Mesostigmata, Opilioacarida+Ixodida and Ixodida) were functionally annotated using eggNOG-mapper version 5^74,82^ (Supplementary file 2.10).

### Hox gene identification and microsynteny analyses

Putative Hox genes were identified from multiple peptide datasets using a reference-guided homology search pipeline. Reference Hox protein sequences from the tick *Ixodes scapularis*, the spider *Parasteatoda tepidariorum*, and the pycnogonid *Pycnogonum litorale* were used as queries in BLASTP searches against species-specific peptide databases, following established approaches for detecting chelicerate Hox gene repertoires and signatures of whole-genome duplication^29,30,80^. Candidate homologs were extracted, combined with reference sequences, aligned using MAFFT, and filtered to remove sequences consisting entirely of gaps. Phylogenetic relationships were inferred using maximum-likelihood analyses in IQ-TREE3 with automated model selection and resulting gene trees were used to identify orthologous and paralogous relationships among Hox genes.

Candidate sequences were further validated through manual inspection and reciprocal BLAST analyses to confirm orthology assignments. The genomic coordinates of all identified Hox genes were extracted from genome annotation files and visualized to assess cluster organization, gene order, transcriptional orientation, and chromosomal distribution. Patterns of Hox cluster fragmentation and genomic rearrangement were compared across Parasitiformes and selected outgroup taxa. To evaluate evidence for whole-genome duplication, Hox gene copy numbers and phylogenetic relationships were examined across all sampled chelicerates, with the pycnogonid *P.litorale* (GCA_964442445.1) included as an outgroup representative. The genomic positions, orientations, and scaffold assignments of identified Hox genes were extracted from genome annotation files and used for plotting using the visualization pipeline (3.hox-plot.py).

### MicroRNA identification and comparative analyses

MicroRNA families were identified using MirMachine version 0.3.0.3^81^ with Chelicerata specified as the target lineage and the protostome covariance model database used for prediction. MirMachine identifies homologs of conserved metazoan and arthropod microRNA families through covariance-model searches against curated MirGeneDB reference profiles. For each species, only filtered high-confidence predictions were retained. The filtered hits output from each MirMachine analysis was used to construct a comparative copy-number matrix to assess lineage-specific expansions and contractions. Conserved developmental microRNAs associated with Hox clusters were examined separately through genomic localization analyses.

### Hox-associated microRNA analyses

The genomic positions of all predicted microRNAs were compared with annotated Hox gene coordinates to identify conserved Hox-associated microRNAs. Particular attention was given to Mir-10, Mir-2, Mir-279, and Iab-4 because of their conserved associations with developmental genes. MicroRNA–Hox positions were visualized across all sampled taxa using a custom script (1.hox-miRNA-map.py and 2.plot-hox-miRNA.py).

### Transposable element analyses

Repeat annotations from RepeatModeler2 were subsequently classified using RepeatMasker into major categories including DNA transposons, LINEs, SINEs, LTR retrotransposons, Simple repeats, Satellites, and Unclassified elements. Repeat abundance was quantified as the proportion of each genome occupied by individual repeat categories. To estimate the relative age of transposable-element insertions, sequence divergence between individual repeat copies and their consensus sequence was calculated using the Kimura two-parameter distance implemented in RepeatMasker version 4.1.866^67^.

To identify lineage-specific patterns of transposable element accumulation and turnover, TE landscapes were generated using RepeatMasker output files by calculating the genomic proportion occupied by repeats from their consensus sequences. Repeat annotations were grouped into major TE classes (DNA transposons, LINEs, SINEs, LTR retrotransposons, Simple repeats, Satellites, and Unclassified elements), with class-specific abundance profiles were normalized by genome size. Low divergence values indicate recent activity and high divergence values indicate older insertions. Divergence distributions were summarized using weighted mean, median, interquartile range, and standard deviation, with repeat lengths used as weights using the custom script (1.repeat-landscapes.py). Smoothed TE landscape curves were visualized for individual genomes and across species to compare the timing and magnitude of historical transposable element activity using the custom script (2.plot_TE-divergence-boxplots.py).

### Genome architecture analyses

Genome architecture was characterized from genome annotation files by extracting exon and intron coordinates for the longest transcript associated with each gene. Gene structural features, including exon length, intron length, gene span, exon number, intron number, and gene compactness, were calculated for each species. Species-level summary statistics were generated and compared across taxa, and distributions of intron lengths, exon lengths, and gene compactness were visualized using the custom script (1.Intron-from-gtf_corr.py).

Gene compactness was calculated using the for each transcript as Total exon length/Gene span, where total exon length corresponded to the cumulative length of all exons within a transcript and gene span corresponded to the genomic distance from the start of the first exon to the end of the last exon. Values approaching 1.0 for highly compact genes containing little or no intronic sequence, whereas lower values indicate increasingly intron-rich gene structures.

## CODE AVAILABILITY

*<u>GenomeCartographer</u>* scripts generated in this study are available at https://github.com/kulkarni-lab/GenomeCartographer/tree/main. The parasitiform linkage group and input empirical data for running the scripts is available at [FigShare link].

## ACKNOWLEDGEMENTS

Authors acknowledge the participation of Apurva Jadhav and V. Y. Deshpande in field work; Gopi Krishnan for discussions on genome assembly trials and Neha Tambe and Nithin K. A. for helping in arranging the supplementary material. We acknowledge the help of A. Sreenivas, Next Generation Sequencing facility in library preparation and sequencing. All analyses were conducted on the Ramanujan High Performance Computing at the CSIR-CCMB, Hyderabad, India. S.K discloses support for this study was from the Ramanujan Fellowship (ANRF/RJF/2023/000045, S.K.), Early Career Research Grant (ANRF/ECRG/2024/000947/LS, S.K.) and OLP0030 (S.K.) grants. A.C. and P.K. were funded by (ANRF/RJF/2023/000045, S.K.) and OLP0030 (S.K.) grants respectively.

## AUTHOR CONTRIBUTIONS

Conceptualization: S.K. Methodology: S.K. Software: S.K., A.C., P.K. Validation: A.C., P.K., D.M. Formal analysis: S.S., A.C., P.K. Investigation: S.S., A.C., P.K., S.K. Resources: S.K. Data curation: A.C., P.K. Writing: All. Writing–Review: All. Visualization: All. Supervision: S.K. Project administration: S.K. Funding: S.K.

## ADDITIONAL INFORMATION

**Supplementary information** The online version contains supplementary materials available at [doi link]

**Correspondence** and requests for materials should be addressed to Siddharth Kulkarni.

