## Supplemental Table for "Ancestral genomic stasis and coordinated genome erosion underlies asymmetric acarine diversification"

### Supplementary files.

1. Synteny

   1.1- Synteny analyses of acarine taxa using ODP.

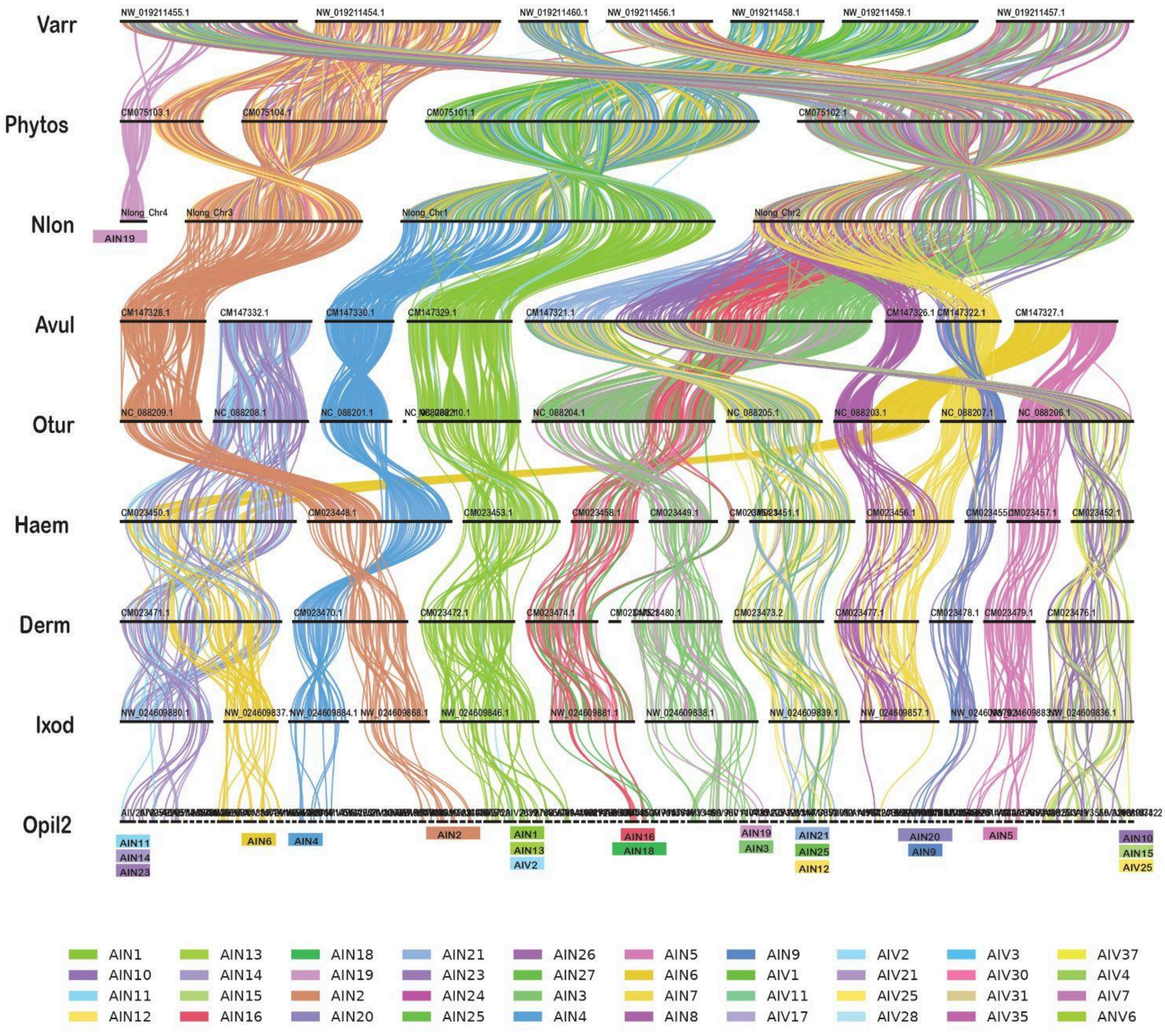


Figure 1.1- A ribbon plot depicting conserved macrosynteny within genomes of *Varroa destructor*, *Phytoseiulus persimilis, Neoseiulus longispinosus, Argas vulgaris, Ornithodoros turicata, Haemaphysalis longicornis, Dermacentor andersoni, Ixodes scapularis* and *Indiacarus* sp. using Parasitiformes linkage groups (ParaLG).

### 1.2- Oxford dot plots (ODPs) of acarine taxa

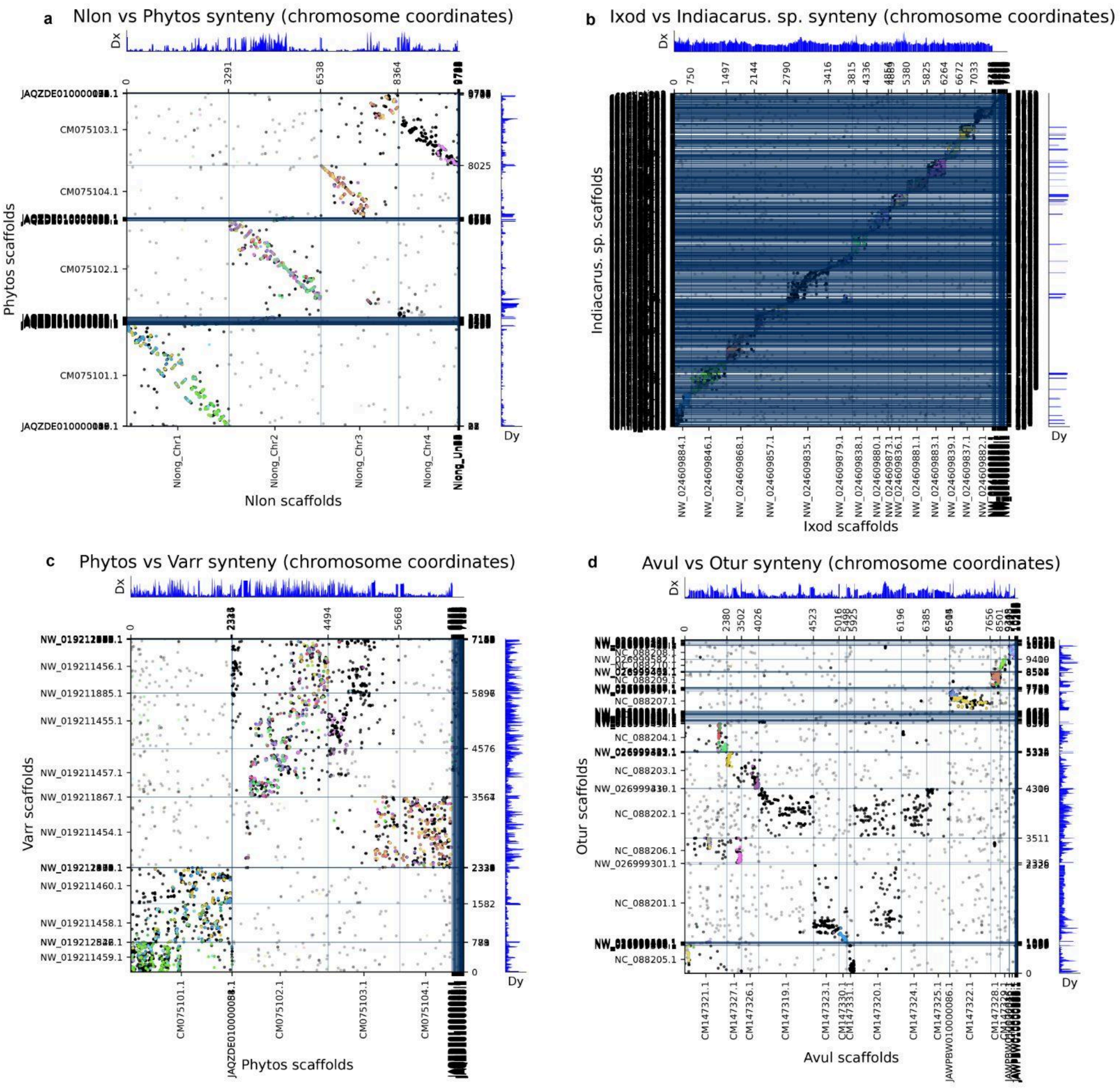


**a.** diamond: 9754 orthologs on 29 Nlon scaffolds and 116 Phytos scaffolds. Colored by protein ids of Nlon in the odp_LGs_from_Avul_Ixod_Nlon_Varr models.
**b.** diamond: 7572 orthologs on 112 Ixod scaffolds and 1948 Indiacarus. sp. scaffolds. Colored by protein ids of Ixod in the odp_LGs_from_Avul_Ixod_Nlon_Varr models.
**c.** diamond: 7164 orthologs on 112 Phytos scaffolds and 46 Varr scaffolds. Colored by protein ids of Varr in the odp_LGs_from_Avul_Ixod_Nlon_Varr models.
**d.** diamond: 10233 orthologs on 73 Avul scaffolds and 97 Otur scaffolds. Colored by protein ids of Avul in the odp_LGs_from_Avul_Ixod_Nlon_Varr models.

**Linkage Group Colours**

| | | | | | | | | | | |
|---|---|---|---|---|---|---|---|---|---|---|
| AIN1 | AIN13 | AIN18 | AIN21 | AIN26 | AIN5 | AIN9 | AIV2 | AIV3 | AIV37 | None |
| AIN10 | AIN14 | AIN19 | AIN23 | AIN27 | AIN6 | AIV1 | AIV21 | AIV30 | AIV4 | p > 0.05 |
| AIN11 | AIN15 | AIN2 | AIN24 | AIN3 | AIN7 | AIV11 | AIV25 | AIV31 | AIV7 | p <= 0.05 |
| AIN12 | AIN16 | AIN20 | AIN25 | AIN4 | AIN8 | AIV17 | AIV28 | AIV35 | ANV6 | |

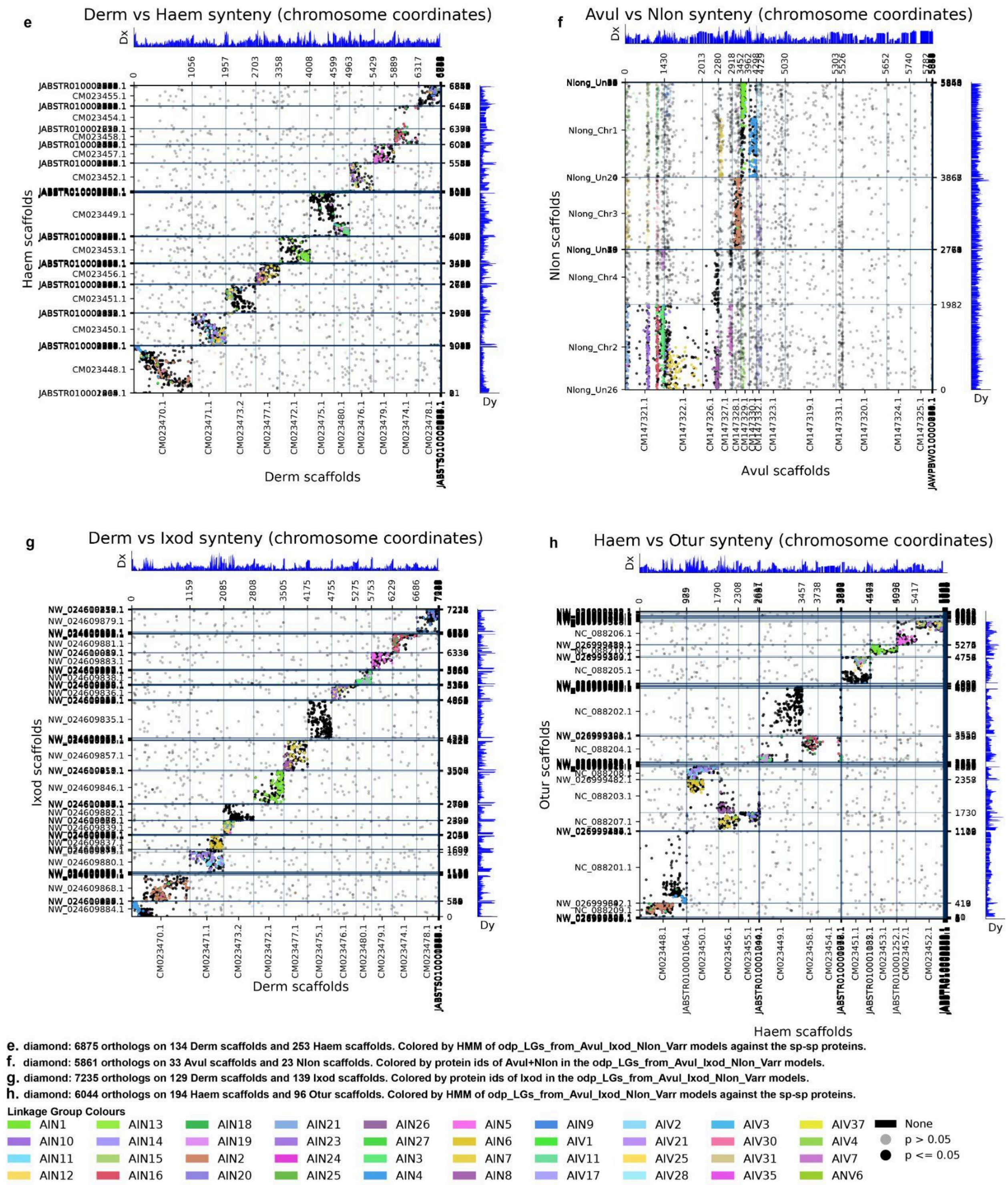


Figure 1.2- Oxford dot plots showing genomic comparison between a) *Neoseiulus longispinosus* and *Phytoseiulus persimilis*, b) *Ixodes scapularis* and *Indiacarus* sp., c) *Phytoseiulus persimilis* and *Varroa destructor*, d) *Argas vulgaris* and *Ornithodoros turicata*, e) *Dermacentor andersoni* and *Haemaphysalis longicornis*, f) *Argas vulgaris* and *Neoseiulus longispinosus*, g) *Dermacentor andersoni* and *Ixodes scapularis*, and h) *Haemaphysalis longicornis* and *Ornithodoros turicata*.

1.3- Self-synteny circos plot of acarine taxa

a

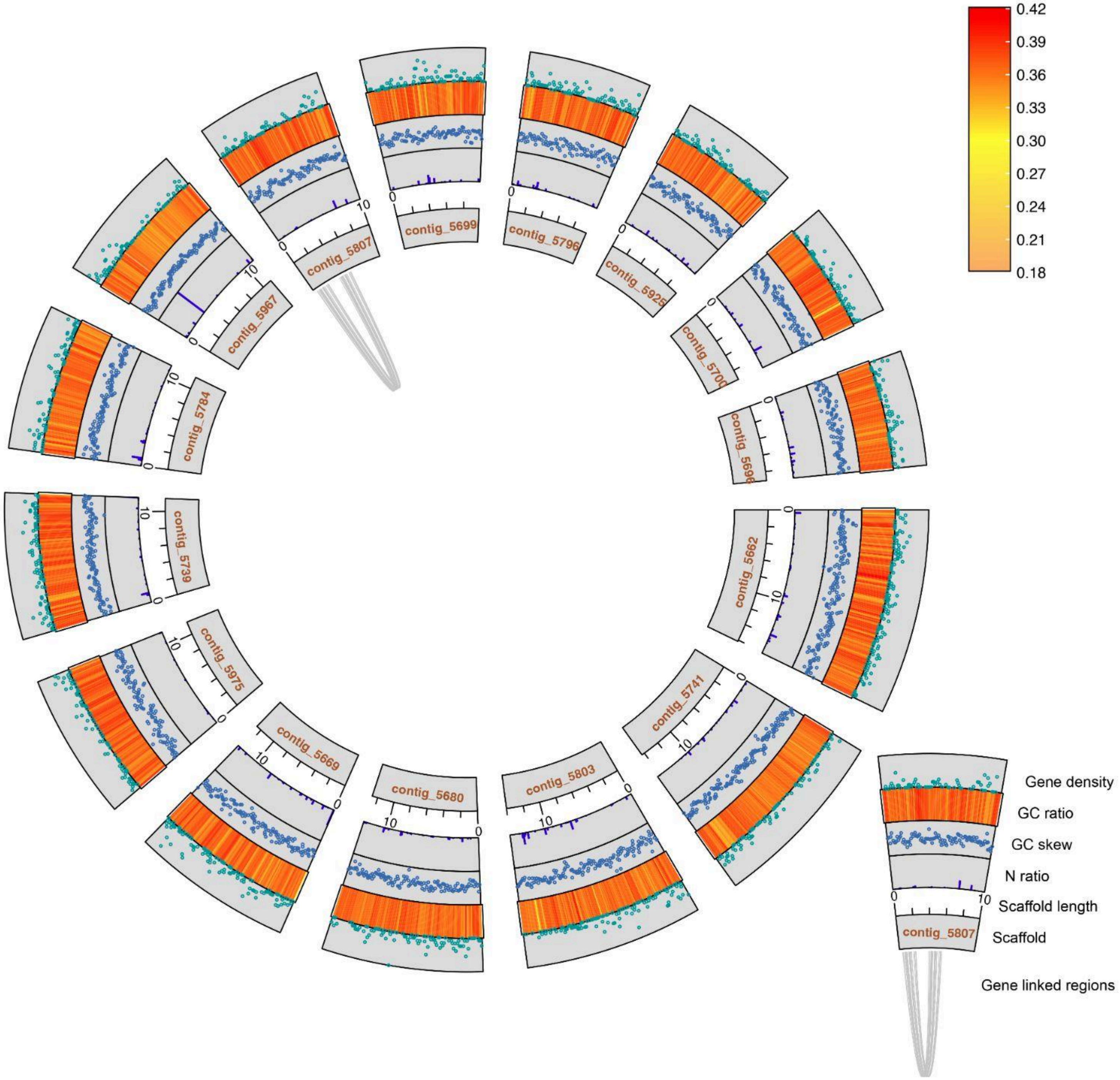

b

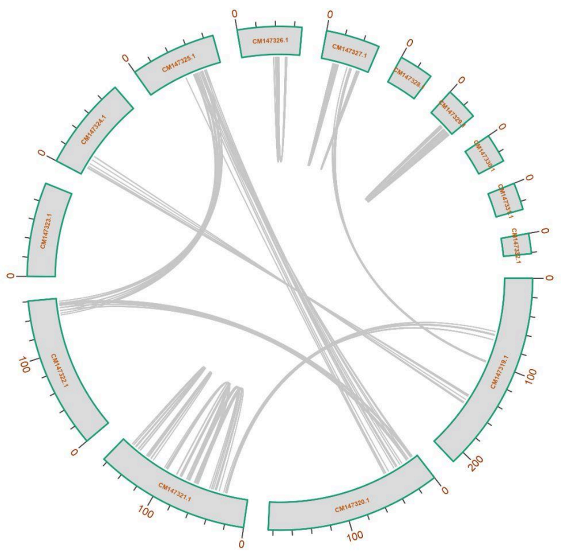

c

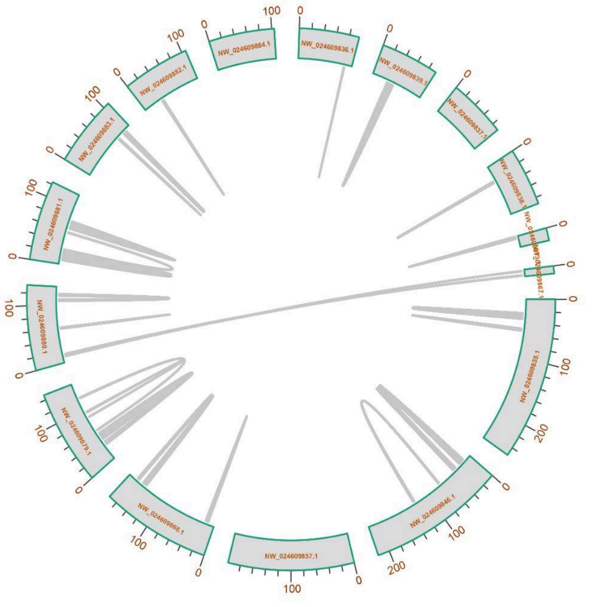

d

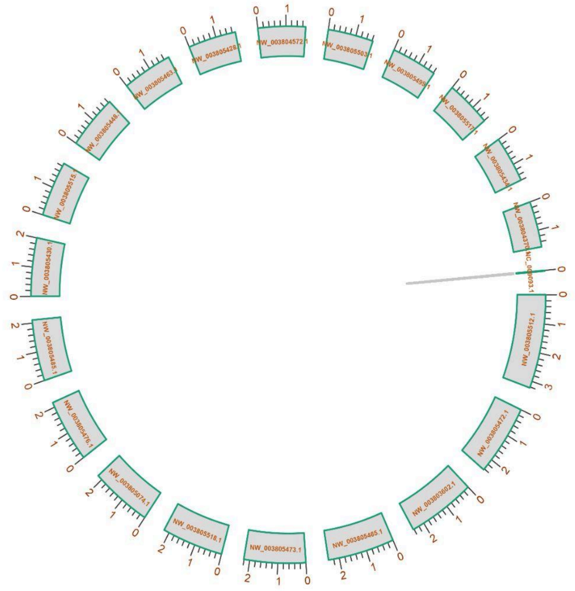

e

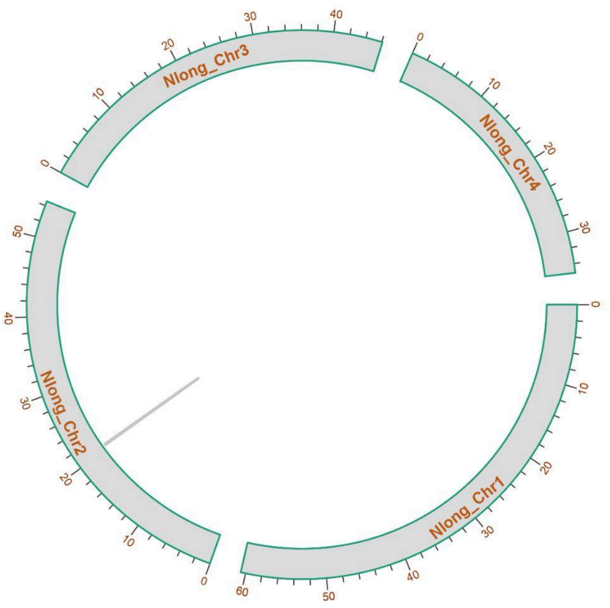
Nlong_Chr3
Nlong_Chr4
Nlong_Chr2
Nlong_Chr1

f

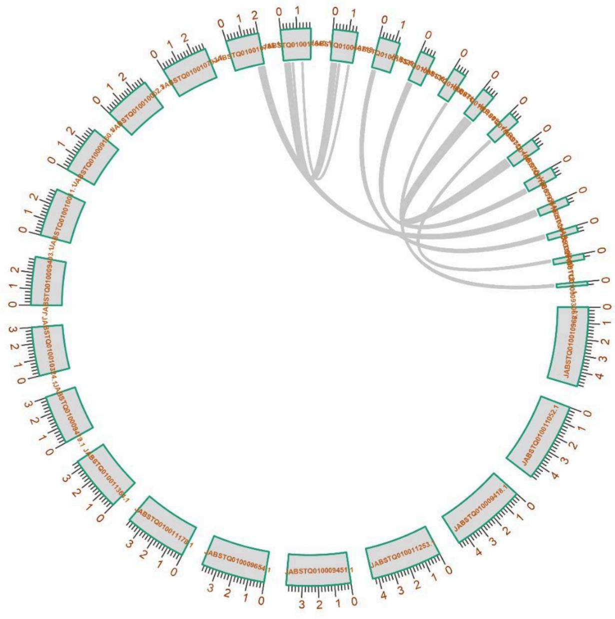

g

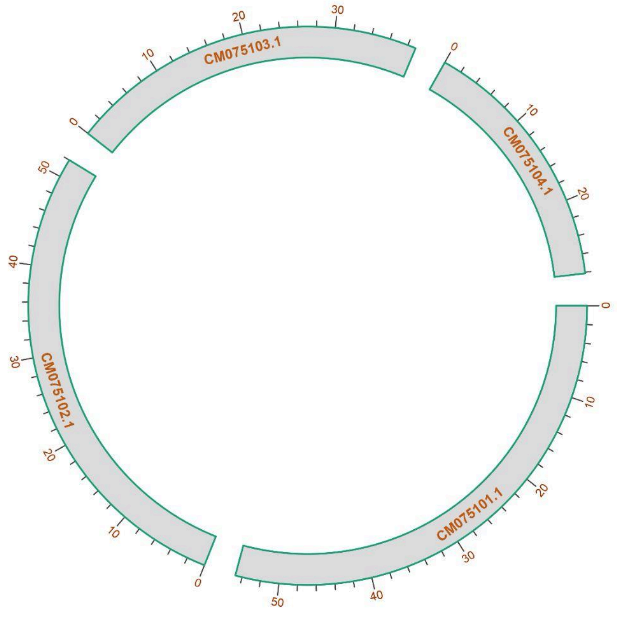
CM075103.1
CM075104.1
CM075102.1
CM075101.1

h

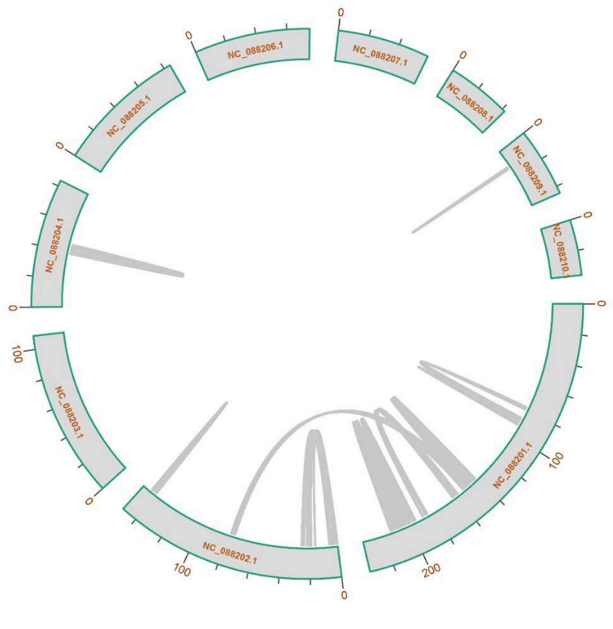
NC_088206.1
NC_088207.1
NC_088205.1
NC_088208.1
NC_088209.1
NC_088204.1
NC_088203.1
NC_088202.1
NC_088201.1

i j

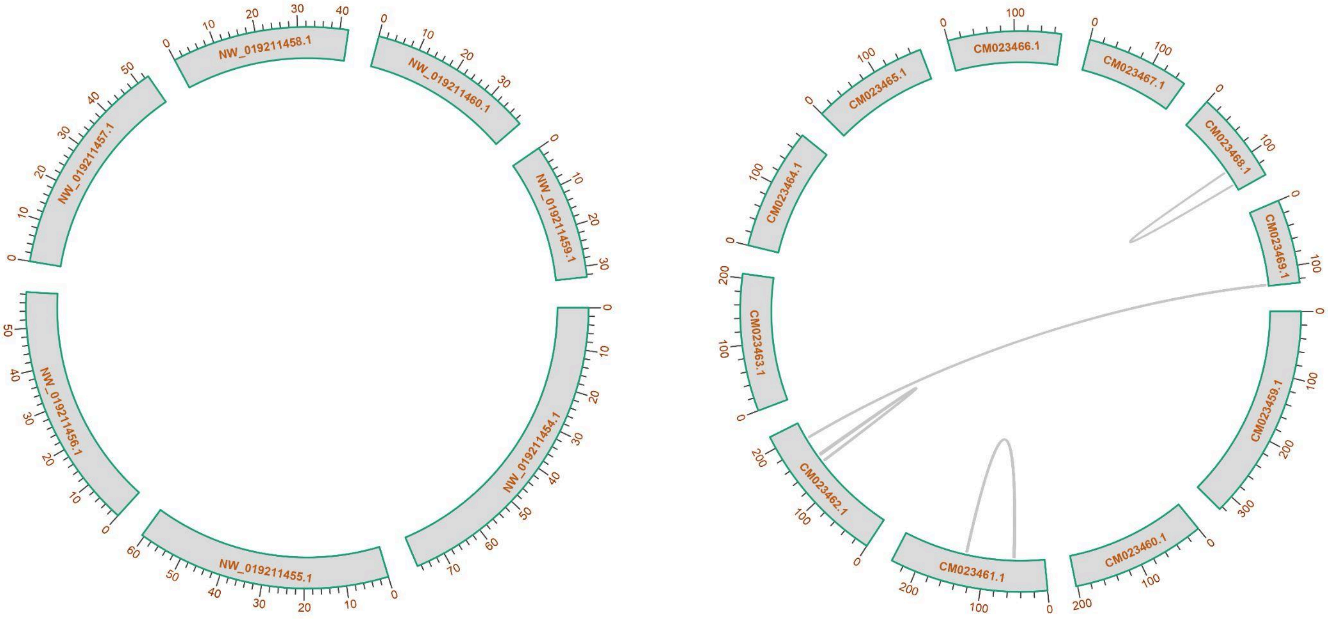


Figure 1.3- Self-synteny plots showing genomic architecture of individual acarine taxa: a) ***Indiacarus* sp.**, b) *Argas vulgaris,* c) *Ixodes scapularis*, d) *Galendromus occidentalis*, e) *Neoseiulus longispinosus*, f) *Ixodes pacificus*, g) *Phytoseiulus persimilis*, h) *Ornithodoros turicata*, i) *Varroa destructor,* and j) *Rhipicephalus microplus***.** (Scaffold length is in Mb)

#### 2. Genomic resources

##### 2.1 Genome assembly of *Indiacarus* sp.

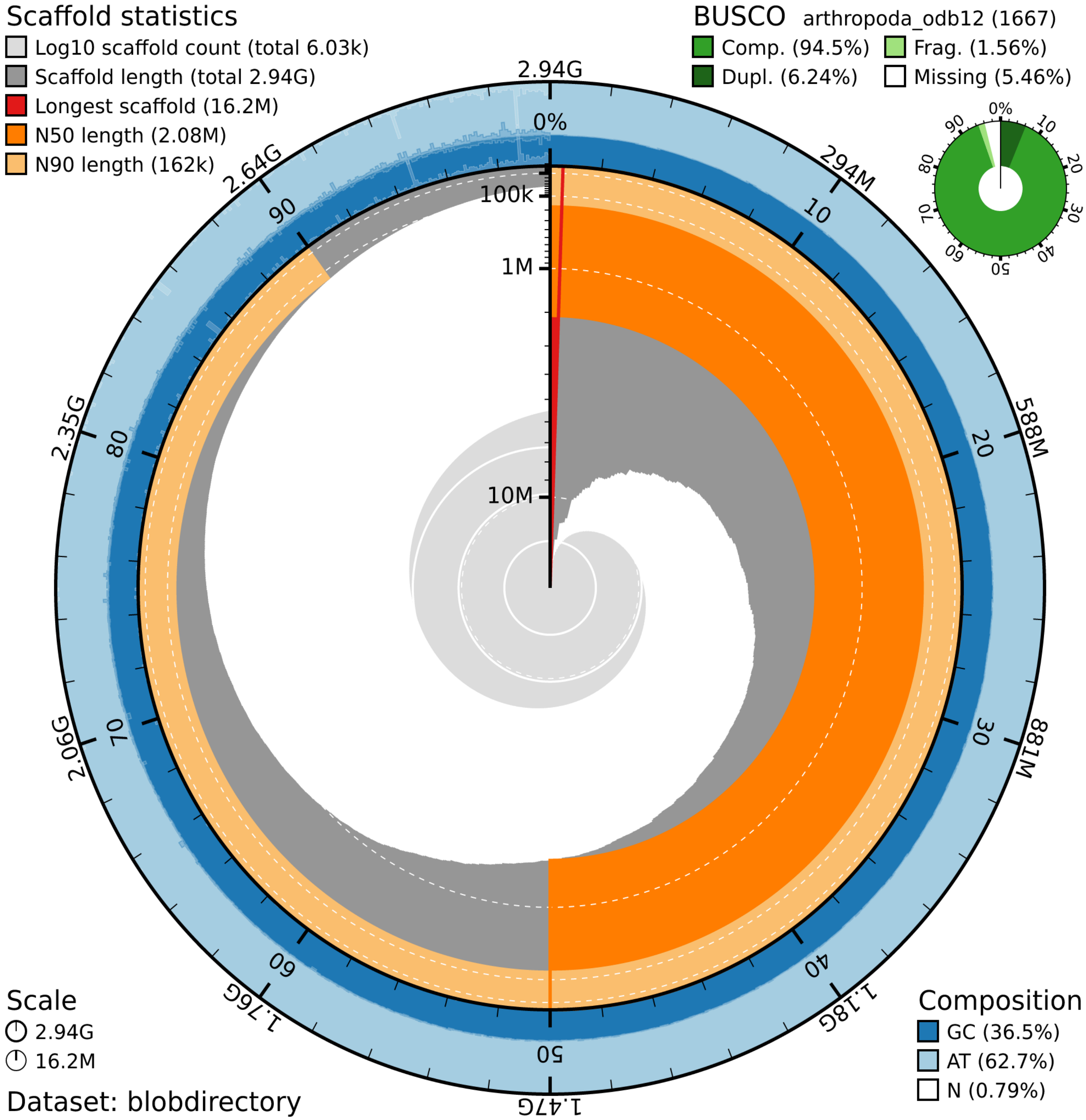


Figure 2.1- Snail plot showing the genome assembly (2.94 Gb) of *Indiacarus* sp. with 94.5% BUSCO completeness, 36.5% GC content, and 2.08 Mb N50 length.

3. Phylogeny

3.1 Dated MCMC phylogeny used for CAFE5 analysis.

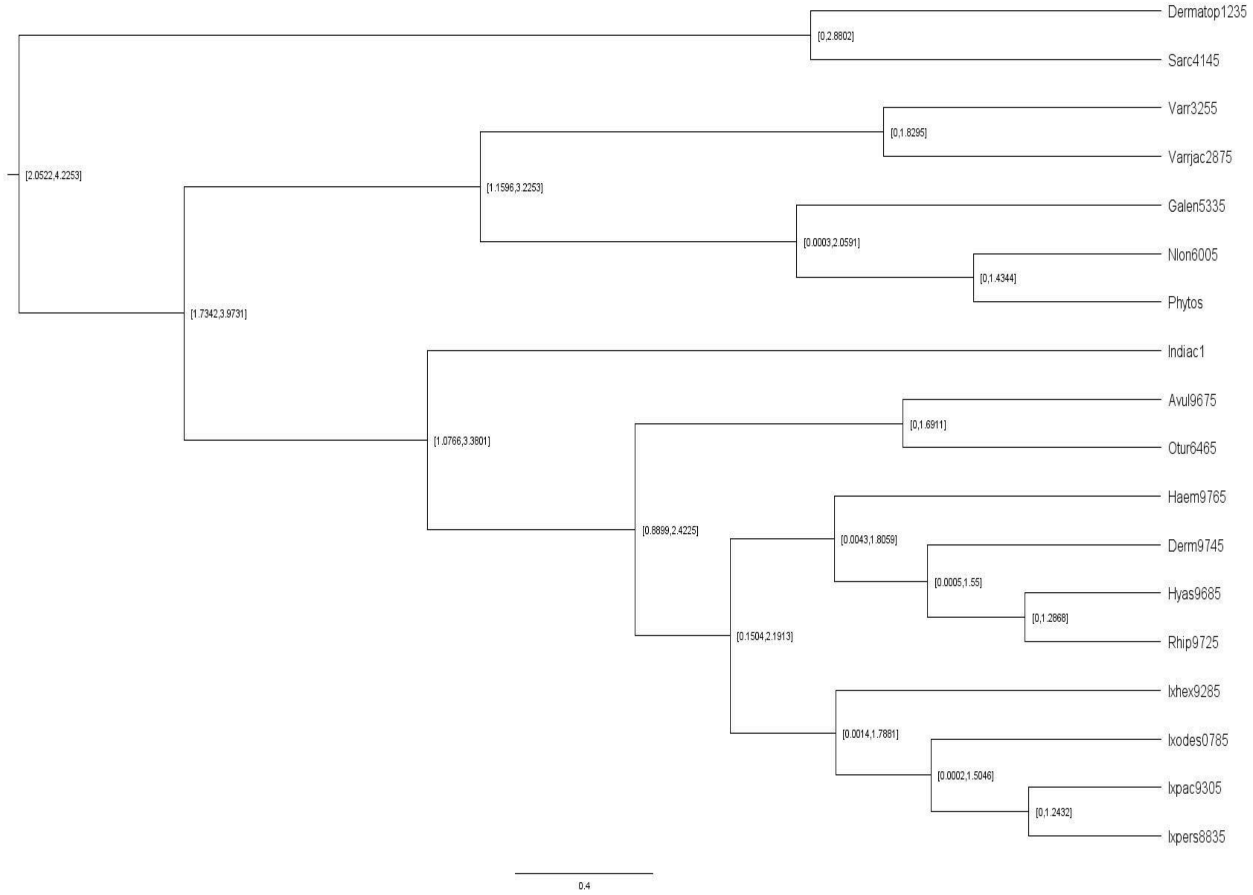

TABLES

2.1- Genome sizes of acarine taxa

| Order | Family | Species Name | Genome size (Mbp) | GenBank Accession |
|---|---|---|---|---|
| Mesostigmata | Phytoseiidae | *Galendromus occidentalis* | 151.460383 | GCA_000255335.2 |
| Mesostigmata | Phytoseiidae | *Neoseiulus longispinosus* | 199.210599 | GCA_052756005.1 |
| Mesostigmata | Phytoseiidae | *Phytoseiulus persimilis* | 181.015949 | GCA_047456425.1 |
| Mesostigmata | Laelapidae | *Varroa destructor* | 368.813097 | GCA_002443255.1 |
| Ixodida | Argasidae | *Argas vulgaris* | 1340.163888 | GCA_055769675.1 |
| Ixodida | Argasidae | *Ornithodoros turicata* | 1071.364447 | GCA_037126465.1 |
| Ixodida | Ixodidae | *Ixodes scapularis* | 2510.615084 | GCA_016920785.2 |
| Ixodida | Ixodidae | *Ixodes persulcatus* | 2444.74277 | GCA_013358835.2 |
| Ixodida | Ixodidae | *Rhipicephalus microplus* | 3265.175345 | GCA_043290135.1 |
| Opilioacarida | Opilioacaridae | *Indiacarus* sp. | 2938.085958 | This study |

2.2- Repeats element proportions (%) in Parasitiform genomes.

| Order | Family | Species Name | Retroelements | SINEs | LINEs | LTR elements | DNA transposons | Unclassified | Small RNA | Satellites | Simple repeats | Low complexity |
|---|---|---|---|---|---|---|---|---|---|---|---|---|
| Mesostigmata | Phytoseiidae | *Galendromus occidentalis* | 3.97 | 0 | 0.75 | 3.21 | 0.97 | 2.62 | 0.04 | 0 | 0.74 | 0.03 |
| Mesostigmata | Phytoseiidae | *Neoseiulus longispinosus* | 0.03 | 0 | 0 | 0.02 | 0.09 | 0 | 0.09 | 0 | 0.88 | 0.06 |
| Mesostigmata | Laelapidae | *Phytoseiulus persimilis* | 6.24 | 0 | 2.12 | 4.12 | 2.55 | 6.93 | 0 | 0 | 0.52 | 0.03 |
| Mesostigmata | Phytoseiidae | *Varroa destructor* | 0.98 | 0.03 | 0.8 | 0.15 | 2.51 | 2.63 | 0.06 | 0 | 1.08 | 0.06 |
| Ixodida | Ixodidae | *Argas vulgaris* | 41.42 | 0.2 | 13.02 | 28.13 | 9.45 | 16.82 | 0.23 | 0.01 | 0.28 | 0.04 |
| Ixodida | Ixodidae | *Ornithodoros turicata* | 24.76 | 0.82 | 8.74 | 14.86 | 6.57 | 32.77 | 0.2 | 0.03 | 0.56 | 0.08 |
| Ixodida | Argasidae | *Ixodes scapularis* | 32.01 | 0.61 | 8.83 | 22.56 | 3.86 | 30.75 | 0.42 | 0.02 | 0.73 | 0.08 |
| Ixodida | Argasidae | *Ixodes persulcatus* | 32.08 | 0.98 | 8.27 | 22.79 | 4.98 | 29.7 | 1.57 | 0.1 | 0.68 | 0.09 |
| Ixodida | Ixodidae | *Rhipicephalus microplus* | 40.48 | 1.87 | 14.66 | 23.95 | 3.37 | 21.31 | 0.68 | 0.06 | 1.25 | 0.04 |
| Opilioacarida | Opilioacaridae | *Indiacarus* sp | 19.85 | 0.19 | 6.11 | 13.53 | 9.88 | 27.97 | 0.84 | 0.04 | 1.53 | 0.15 |

2.3- GC content of taxa used in this study.

| Order | Family | Species Name | GC content (%) |
|---|---|---|---|
| Mesostigmata | Phytoseiidae | *Galendromus occidentalis* | 51.58 |
| Mesostigmata | Phytoseiidae | *Neoseiulus longispinosus* | 50.24 |
| Mesostigmata | Laelapidae | *Phytoseiulus persimilis* | 51.55 |
| Mesostigmata | Phytoseiidae | *Varroa destructor* | 40.92 |
| Ixodida | Ixodidae | *Argas vulgaris* | 48.71 |
| Ixodida | Ixodidae | *Ornithodoros turicata* | 45.82 |
| Ixodida | Argasidae | *Ixodes scapularis* | 46.16 |
| Ixodida | Argasidae | *Ixodes persulcatus* | 46 |
| Ixodida | Ixodidae | *Rhipicephalus microplus* | 45.79 |
| Opilioacarida | Opilioacaridae | *Indiacarus* sp. | 36.79 |

2.4- Gene-family turnover and functions of taxa used in this study.

| Clade | OG | Turnover | Brief functional annotation | Possible phenotype / ecological significance |
|---|---|---|---|---|
| Opilioacarida + Ixodida | OG0000019 | 14 | Acyl-group transferase | Potentially associated with lipid/acyl metabolism and adaptation to specialized diets or host-associated metabolism; parasitism link is uncertain. |
| Within Ixodida | OG0000027 | 10 | Elongation of very-long-chain fatty acids (ELO) | Lipid metabolism; potentially associated with cuticle/surface-lipid specialization and adaptation to prolonged feeding. |
| Within Ixodida | OG0000030 | 10 | Insect cuticle/chitin-binding protein | Cuticle/exoskeletal or barrier-associated function; may relate to ectoparasitic body-surface specialization. |
| Opilioacarida + Ixodida | OG0000040 | 10 | Serpin / leukocyte elastase inhibitor | Strong candidate for host-interaction/immune modulation; particularly relevant to blood feeding and inhibition of host proteases. |
| Opilioacarida + Ixodida | OG0000055 | 12 | Sulfatase-associated protein; extracellular/signaling domains | Possible modification of extracellular substrates and host–parasite interface; could contribute to tissue interaction or nutrient acquisition. |

| | | | | |
|---|---|---|---|---|
| Within Mesostigmata | OG0000057 | −5 | Acetylcholinesterase / carboxylesterase | Potential reduction in detoxification/neurotransmitter-associated esterase repertoire; could reflect ecological specialization. |
| Opilioacarida + Ixodida | OG0000063 | −3 | Arylsulfatase / sulfuric-ester hydrolase | Reduction in extracellular/metabolic sulfatase repertoire; may reflect specialization of substrate utilization. Not clearly parasitism-specific. |
| Within Mesostigmata | OG0000071 | 5 | Serine/threonine protein kinase | Developmental, reproductive and cellular signaling functions; could reflect life-history specialization rather than parasitism specifically. |
| Opilioacarida + Ixodida | OG0000072 | 11 | THAP/zinc-finger, transposase-associated and regulatory domains | Genome regulation/transposable-element activity; may reflect lineage-specific genome remodeling rather than a direct parasitic adaptation. |
| Within Ixodida | OG0000072 | 11 | THAP/zinc-finger, transposase-associated and regulatory domains | Genome/regulatory diversification; potentially contributes to rapid gene-family remodeling in ticks. |
| Within Mesostigmata | OG0000074 | −6 | P450-associated, transposase, ion-channel and LRR domains | Reduced detoxification/regulatory repertoire; potentially consistent with altered chemical exposure in a host-associated niche, but speculative. |
| Opilioacarida + Ixodida | OG0000086 | 10 | Serine carboxypeptidase-like protein | Potential digestive/nutritional adaptation; particularly interesting for blood feeding because carboxypeptidase expansion occurs in blood-feeding mites. |
| Within Ixodida | OG0000088 | 15 | Sensory receptor-associated protein | Potentially important for host detection, environmental sensing and host-finding behavior. |
| Within Mesostigmata | OG0000088 | −5 | Sensory receptor / signaling receptor | Reduced chemosensory repertoire; potentially interesting for reduced environmental searching in host-associated lifestyles. |
| Within Ixodida | OG0000089 | 8 | Zinc-finger/THAP and transposase-associated regulatory proteins | Regulatory/genomic diversification; possible contribution to lineage-specific adaptation. |
| Within Ixodida | OG0000092 | 12 | DNA-binding/regulatory proteins, EGF/VWA/WD-repeat domains | Potentially involved in signaling, extracellular interactions and regulation; host-interface relevance is plausible. |
| Within Ixodida | OG0000095 | 9 | Kunitz, WAP, Kazal, thrombospondin and other protease-inhibitor domains | Strongest parasitism-associated signal: salivary anti-hemostatic/anti-inflammatory activity, immune evasion and blood-feeding adaptation. |
| Within | OG0000102 | 14 | Chitin-binding/denticl | Potential cuticle/barrier specialization; could relate to interactions with |

| | | | | |
|---|---|---|---|---|
| Mesostigmata | | | e-associated protein | hosts or prey, but a parasitic function is not established. |
| Within Mesostigmata | OG0000107 | −4 | Choline dehydrogenase / glycine-betaine metabolism | Reduced osmolyte/metabolic capacity; potentially associated with ecological specialization. |
| Within Mesostigmata | OG0000108 | −5 | Lipid hydrolase / αβ-hydrolase | Reduced lipid-metabolic capacity; potentially relevant to altered nutritional ecology. |
| Within Ixodida | OG0000117 | −3 | P450-associated; mechanosensory/ion-channel and histidine-phosphatase domains | Possible reduction of sensory/metabolic functions; could reflect specialization to a relatively stable host-associated niche. |
| Opilioacarida + Ixodida | OG0000123 | 10 | Calcium-activated chloride channel / epithelial-associated protein | Possible epithelial, osmoregulatory or secretory adaptation; host-interface relevance is plausible but unproven. |
| Within Mesostigmata | OG0000147 | −4 | Heme peroxidase / oxidative-stress response | Potential reduction in oxidative-stress defenses; interesting in relation to host-associated oxidative environments, but not diagnostic of parasitism. |
| Within Mesostigmata | OG0000162 | −4 | Protease-associated, VWF/TIL/mucin/disintegrin domains | Reduction of extracellular/proteolytic host-interface proteins; notably contrasts with tick protease-inhibitor expansions. |
| Opilioacarida + Ixodida | OG0000176 | 9 | Extracellular lectin / pattern-recognition receptor | Innate immune recognition of microbes; potentially important for pathogen exposure in host-associated environments. |
| Within Ixodida | OG0000206 | −3 | Sodium-dependent glucose transporter / MFS transporter | Potential reduction in carbohydrate uptake/transport; potentially consistent with reliance on host-derived nutrients during feeding. |
| Within Mesostigmata | OG0000248 | −5 | ABC transporter / sterol transport | Reduced membrane transport and sterol handling; potentially reflects nutritional specialization. |
| Within Ixodida | OG0000308 | −3 | LIM-domain/zinc-binding protein | Regulatory/cytoskeletal specialization; ecological significance unclear. |
| Within Mesostigmata | OG0000348 | 6 | Serine/threonine protein phosphatase | Cell-cycle, chromosome and developmental regulation; more consistent with cellular/developmental specialization than a specific parasitic phenotype. |
| Within Ixodida | OG0000448 | −4 | PIF1/Helitron-like DNA helicase | Reduction in genome-maintenance/transposable-element-associated functions; likely genome-streamlining rather than direct parasitism. |

FILES

File 2.1- Parasitiformes linkage group designed using *A.vulgaris*, *I.scapularis*, *N.longispinosus* and *V.destructor*.

File 2.2- miRNA copy number of taxa used in this study.

File 2.3- Gene Density of taxa used in this study.

File 2.4- Intron lengths of taxa used in this study.

File 2.5- Gene compactness of taxa used in this study.

File 2.6- Exon lengths of taxa used in this study.

File 2.7- Repeat TE landscape of taxa used in this study.

File 2.8- Canonical HOX-associated miRNAs.

File 2.9- CAFE 5 & time-calibrated phylogeny input.

File 2.10- Gene ontology and EggNOG predictions used in this study.
